# ecDNA amplification of *MYC* drives intratumor copy-number heterogeneity and adaptation to stress in PDAC

**DOI:** 10.1101/2023.09.27.559717

**Authors:** Antonia Malinova, Daniel Schreyer, Elena Fiorini, Davide Pasini, Michele Bevere, Sabrina D’Agosto, Silvia Andreani, Francesca Lupo, Lisa Veghini, Sonia Grimaldi, Serena Pedron, Craig Nourse, Roberto Salvia, Giuseppe Malleo, Andrea Ruzzenente, Alfredo Guglielmi, Michele Milella, Rita T. Lawlor, Claudio Luchini, Christian Pilarsky, Aldo Scarpa, David A. Tuveson, Peter Bailey, Vincenzo Corbo

**Author notes:** These authors contributed equally to this work (Antonia Malinova, Daniel Schreyer, Elena Fiorini). co-last authors (Peter Bailey, Vincenzo Corbo).

## Abstract

Intratumor heterogeneity and phenotypic plasticity drive tumour progression and therapy resistance. Oncogene dosage variation contributes to cell state transitions and phenotypic heterogeneity, thereby providing a substrate for somatic evolution. Nonetheless, the genetic mechanisms underlying phenotypic heterogeneity are still poorly understood. Here, we show that extrachromosomal DNA (ecDNA) is a major source of high-level focal amplification in key oncogenes and a major contributor of *MYC* heterogeneity in pancreatic ductal adenocarcinoma (PDAC). We demonstrate that ecDNA can drive exceptionally high dosage of *MYC* and afford cancer cells rapid adaptation to microenvironmental changes. The continued maintenance of extrachromosomal *MYC* is uniquely ensured by the presence of the selective pressure. We also show that MYC dosage affects cell morphology and dependence of cancer cells on stromal niche factors, with the highest MYC levels correlating with squamous-like phenotypes. Our work provides the first detailed analysis of ecDNAs in PDAC and describes a new genetic mechanism driving *MYC* heterogeneity in PDAC.

## Introduction

Oncogene dosage variation is a major determinant of tumour progression and phenotypic heterogeneity ^1–3^. Focal oncogene amplifications and rearrangements have been demonstrated to underpin oncogene dosage variation and can exist as linear amplifications of contiguous genomic segments or as circular extrachromosomal DNAs (ecDNAs). ecDNAs lack centromeres and therefore segregate unevenly between daughter cells during mitosis ^4,5^. This non-Mendelian pattern of inheritance enables individual cells to accumulate large numbers of ecDNA-bearing oncogenes in response to specific microenvironmental changes ^6^. Rapid depletion of ecDNAs is also observed when cancer cells are no longer exposed to the selective pressure for which they confer enhanced fitness ^6–9^.

Oncogenic amplifications of genes including *GATA6*, *KRAS* and *MYC* have been shown to shape PDAC cancer phenotypes ^3,10–14^. Elevated *MYC* activity promotes biologically aggressive PDAC phenotypes by driving proliferation and remodelling of the tumour microenvironment ^15,16^. *MYC* amplifications are specifically enriched in PDAC liver metastases and are associated with basal-like and squamous subtypes ^15^. Therefore, identifying the genetic events driving transcriptional *MYC* heterogeneity is critical to advance our understanding of disease progression and metastasis. To overcome the limitation of poor neoplastic cellularity of PDAC tissues and enable modelling the dynamics of endogenous oncogene amplifications, we comprehensively characterised a large array of patients derived organoids (PDOs). The integration of PDOs genomes, transcriptomes, and *in situ* analyses with functional studies revealed the role of ecDNA-based *MYC* amplification in driving extensive copy-number heterogeneity and the adaptation of PDAC cells to the depletion of stromal niche factors.

## ecDNAs are a major source of focal oncogene amplifications in PDAC

To characterise the genomic rearrangements that underpin copy number variation in PDAC, we performed whole genome sequencing (WGS) on 41 early passage PDOs established from 39 tumours (**Suppl. Table 1**). The majority of PDOs were established from treatment naive (38/39) and localised tumours. Histopathologically, the majority of PDAC tumours from which PDOs were established displayed a conventional morphological pattern^17^ (36/39), with two containing squamous components (defined as adenosquamous), and one classified as a signet-ring tumour (**Suppl. Table 1**).

Consistent with earlier studies ^18–20^, PDOs exhibited frequent copy-number alterations in canonical PDAC genes including copy number loss of *CDKN2A*, *TP53*, and *SMAD4* and copy number gains in *KRAS*, and *MYC* (**Extended Data** Figure 1a). AmpliconArchitect (AA)^21^ was used to reconstruct genomic regions associated with high copy number gains classifying them as either linear, break-fusion-bridge (BFB), complex, or circular extrachromosomal DNA (ecDNA) amplicons (**Figure 1a** and **Suppl. Table 2**). This analysis revealed that 12 out of 41 PDOs harboured at least one distinct ecDNA (**Figure 1a**). The identification of ecDNAs in PDOs is consistent with earlier whole genome sequencing analyses derived from resected PDAC patient material ^22^. We observed higher fractions of tumour bearing amplifications in PDOs (HCMI) compared to primary tumours (ICGC) (73.17 % vs 66.1 %), including ecDNA amplifications (29.3 % vs 14.2 %) which may be due to increased detection in pure neoplastic populations (**Extended Data** Figure 1b). Importantly, the presence of BFB and/or ecDNA amplifications was associated with poor patients’ outcomes in our cohort (**Extended Data** Figure 1c).

**Figure 1.**
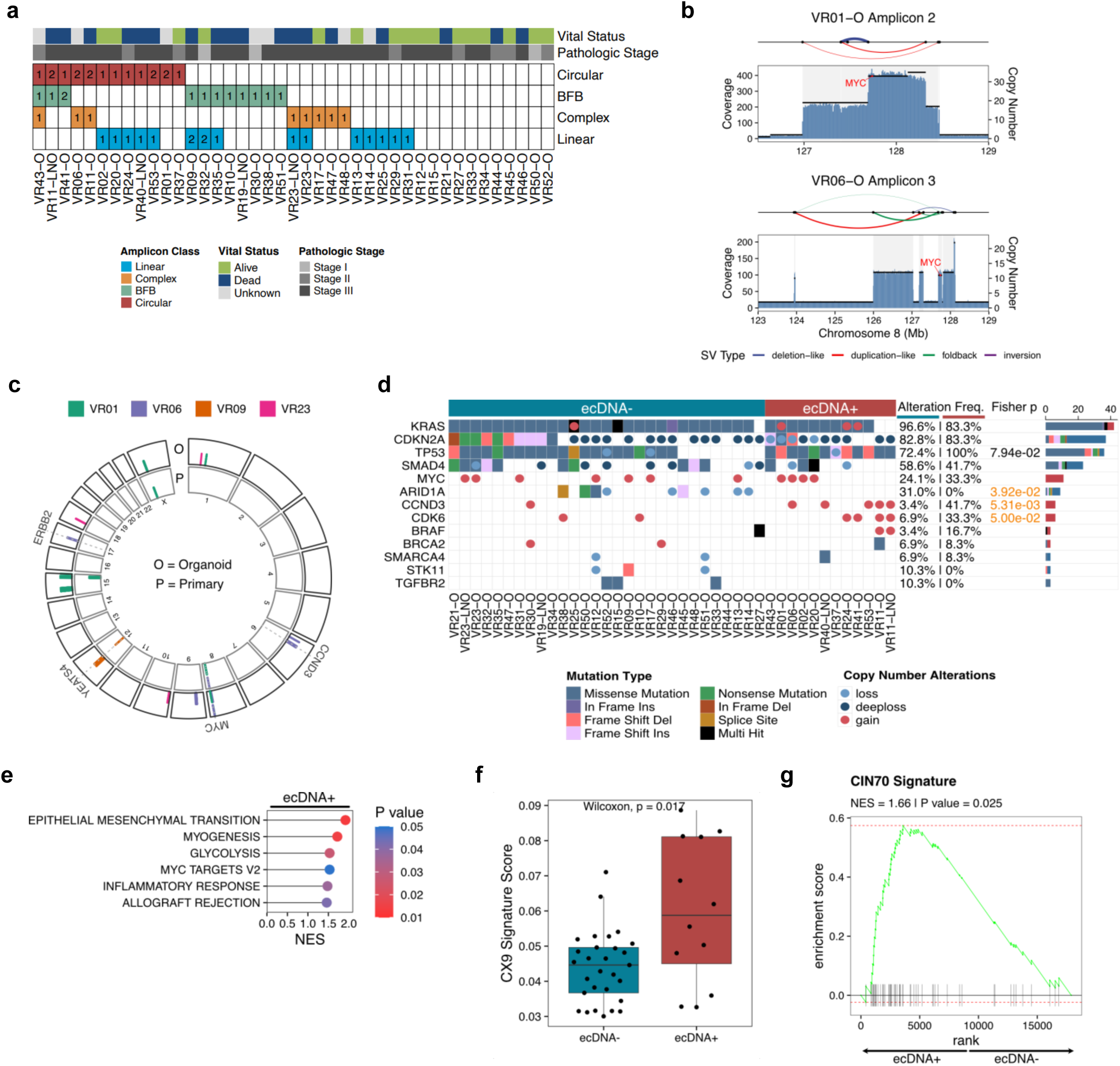
Gene amplification landscape of PDAC. **a**, Amplicon classification based on AmpliconArchitect (AA) of the 41 PDOs subjected to WGS. Number of identified amplicons for each sample is indicated in the plot. Patients pat hol ogi cal st age at time of resect ion and it al status at follow-up is colour-coded. **b**, Genomic view of AA reconstructed amplicon structures spanning the *MYC* locus for the organoids VR01-O and VR06-O with *MYC* on circular amplicons. The genomic view shows coverage depth, copy number segments and structural variant (SV) connections (curves above copy number and coverage view). **c**, Circular plot showing that amplicon regions identified in 4 patients pr ima r t umou r s (P) ar e retained i n t he mat ched organoids (O). **d**, Oncoplot showing the altered genes in 41 PDAC PDOs classified as ecDNA+ (red) and ecDNA-(blue). Mutations were identified using targeted sequencing and copy number calls were derived from the WGS data analysis. The types of alterations are colour– and shape-coded. Gain, copy number >= 3; loss, copy number <= 1, deep loss, copy number <= 0.25. Fishers exact test was utilised to identify associations with genomic alterations in specific genes and ecDNA+ or ecDNA-status. P values below 0.1 are displayed, and significance (p value < 0.05) is highlighted in orange. **e**, Lollipop plot showing gene set enrichment analysis (GSEA) results focused on Hallmark pathways that are significantly enriched among ranked differentially expressed genes of ecDNA+ PDOs (n=7) compared to ecDNA-PDOs (n = 7). **f**, Box plot showing the CX9 chromosomal instability signature enriched in ecDNA+ compared to ecDNA-PDOs. Statistical significance was evaluated using a Wilcoxon rank-sum test. **g**, Gene set enrichment analysis (GSEA) of differentially expressed genes between ecDNA+ (n = 7) and ecDNA-(n = 7) PDOs in the CIN70 transcriptomic signature.

*CCND3* and *MYC* were the most recurrently amplified genes in our cohort of PDOs, while linear amplicons were the most commonly AA-reconstructed amplicon type (**Figure 1a**). Amplifications of *CCND3* were identified in 6 out of 41 PDOs and described as either circular, BFB, or complex amplicons (**Extended Data** Figure 1d). Amplifications of *MYC* were identified in 11 PDOs with 2 PDOs harbouring *MYC* on ecDNA (**Figure 1b**). Circularisation for *in vitro* reporting of cleavage effects by sequencing (CIRCLE-seq)^23^ validated the circular amplicon containing *MYC* in VR01-O (**Extended Data** Figure 2a). *MYC* ecDNAs were derived from contiguous genomic regions on chromosome 8 comprising *MYC* and adjacent genes *PVT1* and *CASC11*. (**Suppl. Table 2**). AA analysis of four primary PDAC samples with matched PDOs demonstrated that the structure of PDO *MYC* ecDNA amplicons between parental tissue and derived PDOs were highly concordant (**Figure 1c, and Extended Data** Figure 2b).

To link patterns of ecDNA gene amplification to key mutational drivers, we performed high-coverage targeted sequencing (**Suppl. Table 3**). PDOs containing ecDNAs displayed bi-allelic inactivation of *TP53* (**Figure 1d**) and were enriched for copy number loss of *CDKN2A* on chromosome 9 and for copy number gains on chromosomes 6 (*CCND3*) and 7 (*CDK6*) (**Figure 1d** and **Extended Data** Figure 2c). Moreover, the presence of an ecDNA was inversely associated with inactivating alterations of the TGFb pathway (**Extended Data** Figure 2d). Whole genome duplications were also more frequently observed in PDOs harbouring an ecDNA (**Extended Data** Figure 2e). Consistent with earlier findings ^24,25^, genes residing on ecDNAs exhibited significantly elevated levels of expression when compared to those associated with other amplicon types (**Extended Data** Figure 2f).

Gene expression programmes commonly linked to biologically aggressive tumours, such as Epithelial-to-Mesenchymal Transition and glycolysis, were significantly enriched in ecDNA PDOs (n = 7) as opposed to non-ecDNA PDOs (n = 7) (**Figure 1e** and **Suppl. Table 4**). ecDNA+ PDOs were also enriched for copy number signatures defining patterns of mid-level amplifications which have been associated with replication stress (CX9, **Figure 1f**) ^26^. Endogenous replication stress can cause genomic instability which in turn may result in ecDNA formation ^27^. Consistent with this idea, ecDNA+ PDOs showed enrichment for a transcriptomic signature (CIN70) of chromosomal instability ^28^ (**Figure 1g**). Overall, we found a heterogeneous landscape of genomic amplifications in PDOs and that ecDNA tumours display features of more biologically aggressive disease.

## ecDNAs are a major source of *MYC* intratumor heterogeneity

To understand how ecDNA might contribute to PDAC intratumor heterogeneity, we focused on *MYC* amplifications representing either extra-(*ecMYC*) or linear intra-chromosomal amplification (*icMYC*). The two high-level amplifications of *MYC* identified in our cohort were predicted to reside on ecDNAs (**Extended Data** Figure 3a). Next, we performed DNA FISH for *MYC* and *Centromere 8 (CEN8*) on metaphase spreads from the 2 *ecMYC* and from 3 *icMYC* PDOs. None of the metaphases from the *icMYC* PDOs (VR02, VR20, and VR23) contained *MYC* positive ecDNAs (**Extended Data** Figure 3b). In contrast, ten to hundreds of individual *MYC* positive ecDNA per nucleus were observed in metaphases prepared from the *ecMYC* PDOs (VR01 and VR06) (**Figure 2a**). Next, we examined interphase nuclei to observe the spatial organisation of FISH positive signals and estimate the cell-to-cell variation for the number of *MYC* copies. In the *ecMYC* PDOs, FISH positive signals tended to cluster in defined regions of the nucleus (**Figure 2b**). Compared to *icMYC* PDOs (**Extended Data** Figure 3b), *ecMYC* PDOs displayed a more heterogeneous distribution of *MYC* amplifications per cell (**Figure 2b**). Significant variability of *MYC* copy-number states was even observed in individual organoids (**Figure 2c**). We then analysed the primary tissues from which the PDOs were established and confirmed the significant cell-to-cell variation of *MYC* FISH foci in the ecDNA bearing tumours (**Figure 2d**). In agreement with existing literature, the presence of ecDNA was associated with higher mRNA levels of *MYC* in the *ecMYC* PDOs **(Extended Data** Figure 3c) and MYC protein expression in paired primary tissues (**Extended Data** Figure 3d). Altogether, our data indicate that ecDNA contribute to extensive copy-number intratumor heterogeneity and drive higher *MYC* expression in PDAC tissues.

**Figure 2.**
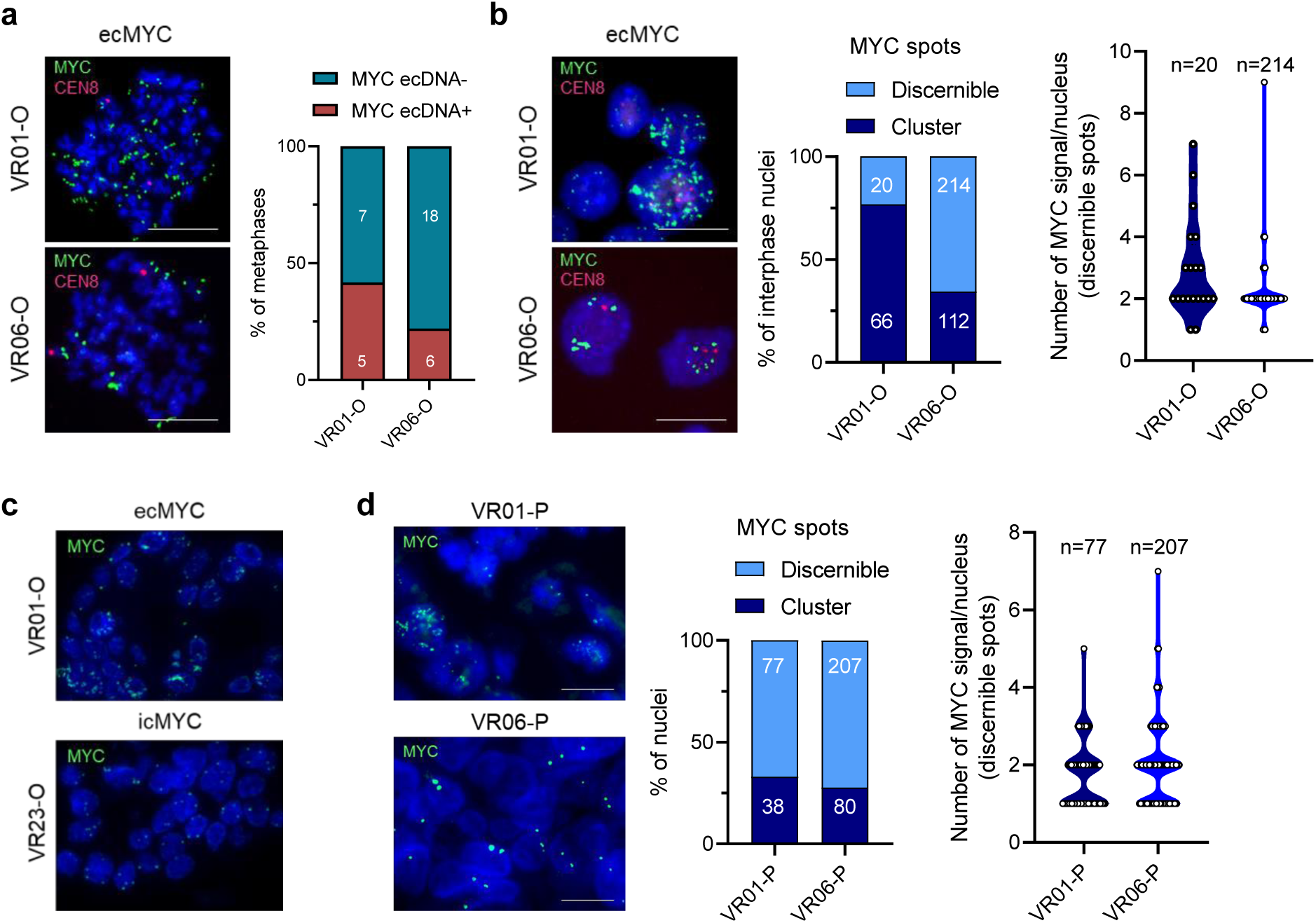
ecDNAs promote intratumor heterogeneity of *MYC* copy number in PDAC. **a**, Representative FISH images validating the presence of *MYC* on ecDNA in VR01 and VR06 PDOs. Scale bar: 20 m (lef t). St acked bar pl ot di spl ai ng t he f requenc of ecDNA+ meta phas es i n VR01-O and VR06-O (right). **b**, Representative FISH images of interphase nuclei of VR01-O and VR06-O. Scale bar: 20 m (lef t). Fr equenc of clust ered *MYC* spots in interphase nuclei (middle). Quantification of discernible *MYC* spots per nucleus is provided on the right. **c**, Representative FISH images of one ec*MYC* (VR01-O, top) and one ic*MYC* (VR23-O, bottom) embedded organoids. **d**, Representative FISH images of embedded VR01 and VR06 primary tissues (P). Scale bar: 20 m (lef t). Fr equenc of clust ered MYC s pot s in pri mar tumo ur s nucl ei (mid dl e). Quantification of discernible *MYC* spots per nucleus is provided on the right.

## ecDNA amplifications of *MYC* drive rapid adaptation to stress

Next, we sought to understand how oncogene-bearing ecDNA dynamically respond to microenvironmental stressors. We took advantage of the well-known dependency of PDAC PDOs on WNT-signalling to impose an artificial selective pressure by removing WNT3A and RSPO ^18,29^ from PDO growth media. EcDNA dynamics were then assessed before and after WNT and RSPO removal (**Figure 3a**). In agreement with previous work ^29^, none of the PDOs tested (n = 9) survived serial passaging in a culture medium lacking both WNT3A and RSPO1 (–WR, **Extended Data** Figure 4a). MYC is a well-established WNT pathway target gene ^30^ and *MYC* expression was rapidly induced in PDOs treated with WNT agonists (**Extended Data** Figure 4b). Strikingly, we found that MYC overexpression was sufficient to bypass the requirement of exogenous WNT3A/RSPO in PDO culture medium (**Extended Data** Figure 4c-d). These results implicated *MYC* as an important driver of WNT-gated survival in PDOs. Therefore, we cultivated *ecMYC* (n = 2), *icMYC* (n = 3), and an individual non-*MYC* amplified PDO in the absence of WR (**Figure 3a**). All the PDOs were established from classical PDAC except for the non-*MYC* amplified PDO (VR09-O), which was derived from a tumour containing signet-ring cells.

**Figure 3.**
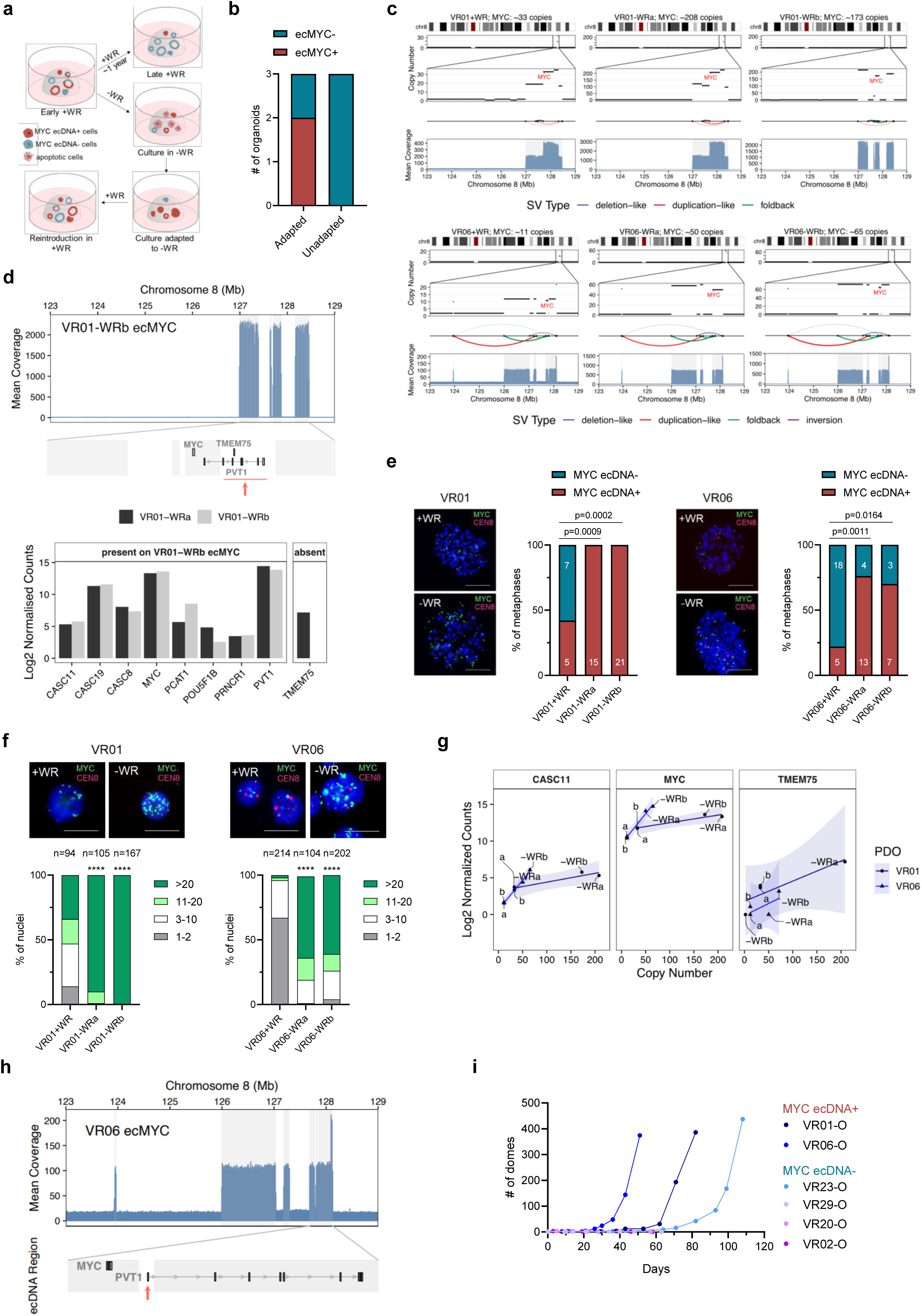
Extrachromosomal *MYC* promotes rapid adaptation to niche factors withdrawal. **a**, Schematic representation of the experimental workflow. **b**, Stacked bar plot displaying proportion of PDOs, classified as *MYC* ecDNA+ or *MYC* ecDNA-, that did or did not adapt to grow in depleted media. **c**, Copy number alterations on chromosome 8 with a focus on *MYC* region, of ecMYC organoids, VR01 (top) and VR06 (bottom), at baseline (+WR) and adapted to depleted media (–WR, 2 biological replicates). SVs that connect amplified regions and form ecDNA and WGS Coverage are displayed below the copy number levels. **d**, Structural difference between putative ecDNAs in VR01-WRa and VR01-WRb (top). The evolved ecDNA structure of VR01-WRb lacks the gene TMEM75. Log2 normalized expression levels of genes that are either present or absent on the ecDNA in VR01-WRb as compared to VR01-WRa (bottom). **e**, Representative FISH metaphases images for VR01 (left) and VR06 (right), at baseline (+WR) and after adaptation to depleted media (–WR). Scale bar: 20 m. Th e st acked bar pl ot s sho t he f requenc of *MYC* ecDNA+ metaphases at baseline and after adaptation (2 biological replicates). P value was calculated using Fisher’s exact test. **f**, Representative FISH interphases images for VR01 (left) and VR06 (right) at baseline (+WR) and after adaptation to depleted media (–WR). Scale bar: 20 µm. Quantification is provided as frequency of nuclei with different ranges of *MYC* spots. ****, p<0.0001 as determined by Chi-square. **g**, Copy number and expression levels of the genes *MYC*, *CASC11*, and *TMEM75*, in VR01 and VR06 at baseline (a, b) and after adaptation (–WRa, –WRb). **h**, Genomic view of VR06 ecMYC segments (highlighted in grey) and the location of *MYC* and *PVT1*. The *PVT1* starting region is absent on the VR06 ecMYC. **i**, Growth curve of MYC ecDNA+ (n = 2) and MYC ecDNA-(n=4) organoids in –WR media. Culture growth is represented as number of domes (50 µl Matrigel/dome).

The withdrawal of WR from the medium led to the rapid extinction of 3 cultures, including two with low-level copy number gains of *MYC* (VR02 and VR20). Conversely, the two *ecMYC* and one *icMYC* PDOs consistently adapted (aPDOs) to the niche-factors depleted conditions (**Figure 3b**). Then, we performed a detailed molecular characterisation of both parental and adapted PDOs. To minimise the confounding effect of the organoid culture medium ^31^, both parental and adapted PDOs were cultivated in the same medium lacking WR and TGF-b/BMP inhibitors (A83-01 and Noggin) before RNA-seq analysis and immunophenotyping. The integrated analysis of genomic and transcriptomic data excluded that PDOs adaptation to WR withdrawal was associated with alternative activation of the WNT pathway (**Extended Data** Figure 4e-i, and **Suppl. Table 5).** Previous work has established that PDOs can create their own niche through the endogenous production of Wnt ligands ^29^. To further confirm niche independence for the adapted PDOs, we treated PDOs with the porcupine inhibitor C59, which blocks endogenous production of biologically active Wnt ligands. In keeping with the niche independence, the adapted PDOs were completely insensitive to porcupine inhibition (**Extended Data** Figure 4j).

To determine how the microenvironmental stress affected chromosomal and extrachromosomal *MYC* dynamics, we initially applied AA to WGS from two adapted cultures from each PDOs (**Figure 3c** **and Extended Data** Figure 5a). EcDNA containing *MYC* persisted in adapted PDOs, increased their integer copy number, and in one instance evolved its structure. No evidence for circular amplicons was found in VR23-O (*icMYC* PDO) (**Extended Data** Figure 5a). The AA-reconstructed circular amplicons for the adapted VR06-O were highly concordant with the circular amplicon described for the parental culture. Compared to the circular amplicon described in the parental culture and persisting in one of the aPDO, an individual genomic locus (*TMEM75*) was not included in the ecDNA structure described in VR01-WRb (**Figure 3d**). Accordingly, RNA-seq did not detect expression of *TMEM75* in the corresponding aPDO (**Figure 3d**).

The accumulation of *MYC*-containing ecDNA in aPDOs was confirmed by FISH on metaphase spreads. Every metaphase and almost all the metaphases from the adapted VR01-O and VR06-O, respectively, exhibited copy number accumulation of ecDNAs (**Figure 3e**). The adaptation was also associated with a dramatic increase in the mean *MYC* copy-number and per cell-distribution which indicates that the ecDNA is under positive selection (**Figure 3f**). In the *icMYC* PDO, adaptation was associated with either a mild increase in *MYC* copy-number (VR23-WRa) or polysomy of chromosome 8 (VR23-WRb) (**Extended Data** Figure 5b) in the absence of changes in ploidy (**Extended Data** Figure 5c). Upregulation of *MYC* at mRNA and protein levels in adapted PDOs was consistent with increased *MYC* copy-number and was more dramatic in *ecMYC* than *icMYC* PDO (**Extended Data** Figure 5d-e).

We then compared RNA-seq data from parental and adapted *ecMYC* PDOs to gain further insights into the transcriptional changes associated with the accumulation of ecDNAs. For almost all genes on the ecDNA amplicons, the increased mRNA expression in adapted organoids mirrored the increased ecDNA copy-number (**Figure 3g**). However, the effect size for the increase of *MYC* and other genes between the two cultures was not always consistent with the predicted copy-number changes, suggesting additional regulatory mechanisms controlling gene expression. A known tumour suppressor element that acts i*n cis* to reduce *MYC* expression is the promoter of the long-noncoding RNA *PVT1* ^32^. The AA-predicted structure for the *MYC* ecDNA in VR06-O lacked the promoter and the first exons of PVT1, thereby providing an explanation for the higher level of *MYC* in VR06-O as compared to VR01-O (**Figure 3h**). Based on our results, we postulated that *MYC* on ecDNA would enable more rapid adaptation than possible through chromosomal inheritance. Therefore, we cultivated parental PDOs in WR depleted media and monitored the dynamics of niche independency acquisition. Consistent with the rapid accumulation of ecDNA and the increase in *MYC* output, the adaptation to the imposed stress was more rapid for VR06 (**Figure 3i**).

## Maintenance of ec*MYC* in PDAC organoids

EcDNAs can be lost by cancer cells in the absence of a selective pressure or upon microenvironmental changes (i.e., drug treatment) that impose negative fitness ^6–9^. Elimination of ecDNA can occur through the integration of an ecDNA into a chromosomal location to form a homogeneously staining region (HSR)^8,33,34^, which are considered as a latent reservoir of ecDNAs. Therefore, we assessed whether ecDNAs are selectively maintained by reversing the selective pressure imposed on PDOs. We reintroduced WR in the culture medium of aPDOs and, in parallel, performed long-term passaging (over 1 year of continuous culture) of parental PDOs in standard medium. After few passages (∼5) in a medium supplemented with WR, we observed a rapid decrease in the number of metaphases containing *MYC* on ecDNA and accordingly of the mean *MYC* copy-number and per cell distribution in aPDOs (**Figure 4a and b**). Following extensive cultivation of parental PDOs in standard organoid medium, *MYC* containing ecDNAs were lost (VR06-O) or substantially reduced (VR01-O) (**Figure 4c and d**). The reduction of ecDNA in VR01-O was associated with the emergence of HSR (Figure 4C), which were instead not observed in VR06-O. Accordingly, the reintroduction of the selection pressure (i.e., WR withdrawal) to the high-passage VR06-O cultures led to rapid extinction of the culture after few weeks with no evidence for the generation of ecDNA containing *MYC* (**Figure 4e**). Altogether, our results suggest that the loss of the selective pressure can have different outputs on ecDNA-containing cells.

**Figure 4.**
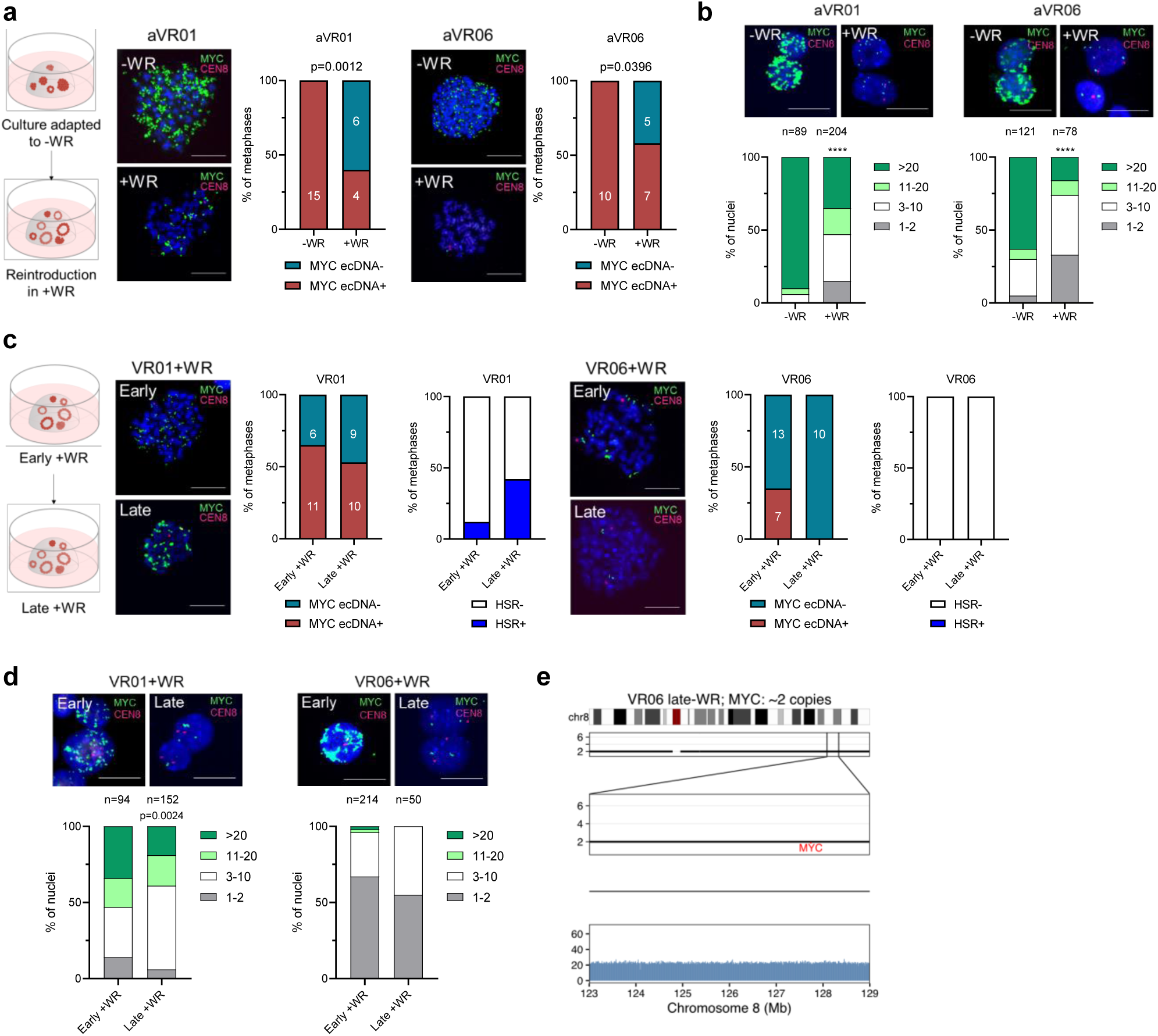
Depletion of ecDNA from PDOs culture upon removal of selection pressure. **a**, Representative FISH metaphase for adapted VR01 and VR06 (aVR01 and aVR06) cultured in +WR media for 5 passages, –WR condition is used as control. Scale bar: 20 µm. Quantification is provided as changes in frequency of *MYC* ecDNA+ metaphases. Significance was assessed by Fisher’s exact test. **b**, Representative FISH interphases for aVR01 and aVR06 cultured in +WR media for 5 passages, –WR condition is used as control. Scale bar: 20 µm. Quantification is provided as frequency of nuclei with different range of *MYC* spots. ****, p<0.0001 by Chi-square. **c**, Representative FISH metaphase for VR01-O and VR06-O cultured in +WR at early and late passages. Scale bar: 20 µm. Quantification is provided as changes in frequency of *MYC* ecDNA+ and HSR+ metaphases. **d**, Representative FISH interphases for VR01-O and VR06-O cultured in +WR at early and late passages. Scale bar: 20 µm. Quantification is provided as frequency of nuclei with different range of MYC spots. Significance was assessed by Chi-square. **e**, Copy number alterations on chromosome 8 (with a focus on *MYC* region) of VR06 late passage after few passages in depleted media (–WR). WGS Coverage is displayed below the copy number level.

## ecDNAs and cell phenotypes in PDAC

Both immunophenotyping (**Extended Data** Figure 6a) and pathway analysis from RNA-seq data demonstrated reduced proliferation index for PDOs with accumulation of ecDNAs (**Extended Data** Figure 6b). Together with the observation that ecDNA are spontaneously lost in the absence of or upon neutralisation of the selection pressure, our results suggest a fitness cost for the maintenance of *MYC* on ecDNA in PDOs. EcDNA-driven cancer cells have been shown to display increased levels of phosphorylated histone H2AX ^27^, which is required for the assembly of DNA damage response as well as for the activation of checkpoint proteins which might arrest cell cycle progression ^35^. Moreover, anti-neoplastic treatments known to activate the DNA damage sensing machinery have been shown to promote the loss of ecDNA through as yet uncharacterised mechanisms ^36–38^. In PDOs, the levels of phosphorylated H2AX (gH2AX) was positively correlated with ecDNA copy-number but not with *MYC* levels (**Figure 5a**). Accordingly, the reduction of ecDNA copies due to the removal of the imposed artificial selection was associated with reduced levels of gH2AX (**Extended Data** Figure 6c). In a heterogeneous organoid culture, MYC protein expression might serve as a proxy for *ecMYC* copy number. We found that gH2AX foci were particularly prominent in *MYC*^high^ nuclei and *MYC* expression positively correlated with the intensity of gH2AX staining (**Figure 5b**). Nonetheless, neither the forced overexpression of *MYC* through a lentiviral vector nor the adaptation of a non *ecMYC* PDOs induced elevation of gH2AX in PDOs (**Extended Data** Figure 6d-e). EcDNA copy-number also correlated with increased levels of the apoptotic marker cleaved PARP (cPARP) (**Figure 5a**), thereby suggesting that accumulation of ecDNA might not be beneficial for cancer cells unless providing a survival advantage.

**Figure 5.**
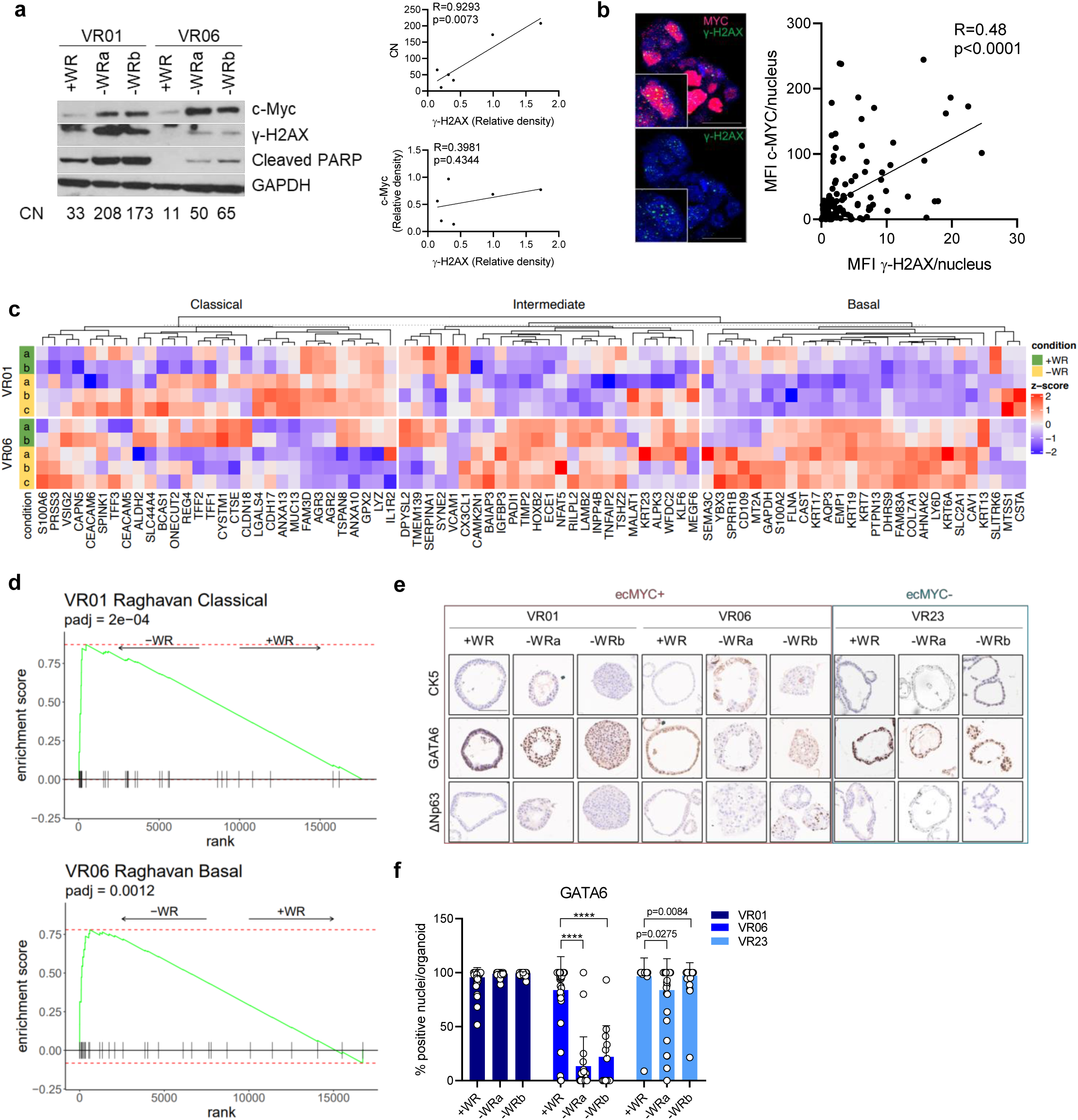
*MYC* levels and neoplastic cell states. **a**, Immunoblot analysis in whole cell lysate of ec*MYC* organoids at baseline and after adaptation to –WR. GAPDH was used as loading control. CN: WGS-based copy number (left). Scatter plot showing the correlation of γ-H2AX protein level and copy number (CN) (top) or c-Myc protein level (bottom). **b**, Representative confocal microscopy images of anti-c-Myc and anti-y-H2AX immunofluorescence in one ec*MYC* organoid at baseline. Scale bar: 20 µm. Pearson R coefficient was used to calculate the correlation between the mean fluorescent intensity (MFI) of c-Myc and γ-H2AX. **c**, Heatmap displaying the expression of Classical, Intermediate, and Basal genes from Raghavan et al.^31^ in baseline (2 biological replicas) and adapted (3 biological replicas) ec*MYC* organoids. **d**, Gene set enrichment analysis (GSEA). Top panel, enrichment of Classical geneset computed over the ranked lists of VR01 differentially expressed genes, derived from the comparison of –WR and +WR samples. Bottom panel enrichment of Basal geneset computed over the ranked lists of VR06 differentially expressed genes, derived from the comparison of –WR and +WR samples. **e**, Representative immunohistochemistry for cytokeratin 5 (CK5), GATA6, and ΔNp63 of parental (+WR) and adapted (–WR) organoids. Scale bar: 100 µm. Quantification for GATA6 is provided in **f** as frequency of GATA6+ nuclei per organoid, at least 20 organoids were analysed for each condition. P values were calculated by Two-way ANOVA. ****, p<0.0001.

We then sought to assess the phenotypic consequences of ecDNA accumulation in PDAC organoids. The aPDOs bearing *ecMYC* and displaying the highest *MYC* expression showed marked morphological changes (**Extended Data** Figure 7a). As opposed to the *icMYC* PDOs (VR23), the two *ecMYC* PDOs lost their characteristic cystic-like structure to display a solid growth pattern with cytological sign of less differentiated tumours (**Extended Data** Figure 7a). In agreement with previous studies ^39,40^, targeting of *MYC* transcription with 500nM of the BRD4 inhibitor JQ1 ^41^ dramatically reduced *MYC*-ecDNA interphase hubs (**Extended Data** Figure 7b) and lowered the level of *MYC* mRNA (**Extended Data** Figure 7c) in *ecMYC* aPDOs. Conversely, levels of *MYC* were almost unaffected by the JQ1 treatment in the *icMYC* aPDOs (**Extended Data** Figure 7c). Furthermore, JQ1 preferentially reduced cell viability of *ecMYC* PDOs over *icMYC* PDO (**Extended Data** Figure 7d). Elevated *MYC* expression and MYC related gene programs are significantly enriched in tumours with basal-like/squamous identity ^10,15^. Therefore, we evaluated whether *ecMYC* accumulation affected the cell states of the adapted PDOs. The accumulation of *ecMYC* was not associated with shifts in PDO cell states (**Figure 5c**). The accumulation of *ecMYC* rather strengthened the classical and the basal programs ^31,42^ in VR01-O and VR06-O, respectively (**Figure 5d**). Conversely, adaptation of *icMYC* PDO to WR withdrawal was associated with more dramatic changes in cell states (**Extended Data** Figure 7e). Changes in transcriptional cell states were concordant with immunophenotypic data, with the PDOs displaying the highest *MYC* dosage (VR06) showing expression, although heterogeneous, of squamous markers (CK5 and DNp63) and reduction of the classical marker GATA6 compared to the parental culture (**Figure 5e and f**).

## Discussion

Intratumor heterogeneity and phenotypic plasticity drive tumour progression and therapy resistance. Oncogene dosage variation contributes to cell state transition and phenotypic heterogeneity ^1–3^, thereby providing a substrate for somatic evolution. Nonetheless, the genetic mechanisms underlying phenotypic heterogeneity are still poorly understood. While the transcriptional output of an oncogene can be specified by either genetic or non-genetic mechanisms, oncogenic activation is often driven by focal amplifications ^43^. ecDNAs are emerging as important mediators of intratumor heterogeneity and therapy resistance in cancer ^6^. Thousands of ecDNA copies may accumulate in a cancer cell and supercharge oncogene expression due to increased chromatin accessibility and enhancer hijacking ^25,39^.

In PDAC, the emergence of copy number amplifications in oncogenes, such as *GATA6*, *KRAS* and *MYC*, defines the evolutionary trajectory of the tumour ^3,44^. Sustained *MYC* activity is required for maintenance and progression of PDAC ^15,16,45^. Elevated *MYC* expression defines a subset of PDAC cells with high metastatic capability and amplifications of MYC are specifically enriched in metastatic PDAC ^15,16^.

Here, we provide the first detailed analysis of ecDNAs in PDAC. We demonstrate that ecDNAs are a major source of high-level amplifications in key PDAC oncogenes and a major contributor of *MYC* heterogeneity in PDAC. We observed different mechanisms of *MYC* amplification. PDOs and tissues harbouring *MYC* on ecDNA displayed significant heterogeneity of *MYC* copy number and higher *MYC* expression compared with tumours having *MYC* on chromosomal DNA. Nonetheless, we found that the transcriptional output of an oncogene from ecDNA cannot be simply inferred from the ecDNA copy-number and *cis*-regulatory elements on the amplicon (e.g., PVT1 promoter) need to be considered. The combined analysis of tissues and PDOs suggest that the generation of ecDNA is likely a late event in the history of pancreatic cancer.

EcDNA were reported in tumours displaying genomic and transcriptomic features of advanced diseases. In line with previous reports^27^, ecDNA in PDOs correlated with the levels of gH2AX, a well-established marker of DNA damage and mitotic checkpoint activation ^35^. In agreement with that, the accumulation of ecDNAs, but not oncogene levels *per se*, was associated with abundant gH2AX foci, reduced proliferative index, and increased apoptotic cell death. Our result suggests that the large number of ecDNA might not be tolerated unless providing enhanced fitness in specific microenvironmental conditions.

Accordingly, the removal of the selection pressure was associated with the rapid reduction of ecDNA elements and of the levels of gH2AX. Similarly, the extensive cultivation of PDOs in the absence of a selection pressure led to either the incorporation of ecDNA into HSR or the irreversible loss of ecDNA.

Mimicking the depletion of stromal niche factors ^18,29^, we show that ecDNAs represent important genomic adaptations that endow tumour cells with the ability to rapidly elevate oncogene expression in response to microenvironmental stressors. Our data further suggest that elevated *MYC* activity is fundamental to the acquisition of stromal independence in pancreatic cancer. While showing that ecDNA is a major source of *MYC* expression heterogeneity in PDAC, we did not conclusively demonstrate the impact of elevated *MYC* levels on neoplastic cell states. Elevation of *MYC* expression due to ecDNA affected PDOs morphology, created cancer cell addiction to transcriptional *MYC* output, but did not induce a molecular class switch. Additional cell intrinsic or cell extrinsic factors might be crucial to the definition of neoplastic cell states. Therefore, further studies will need to address the interaction between ecDNA and cell extrinsic inputs.

## Figure Legends

**Extended Data Fig 1.**
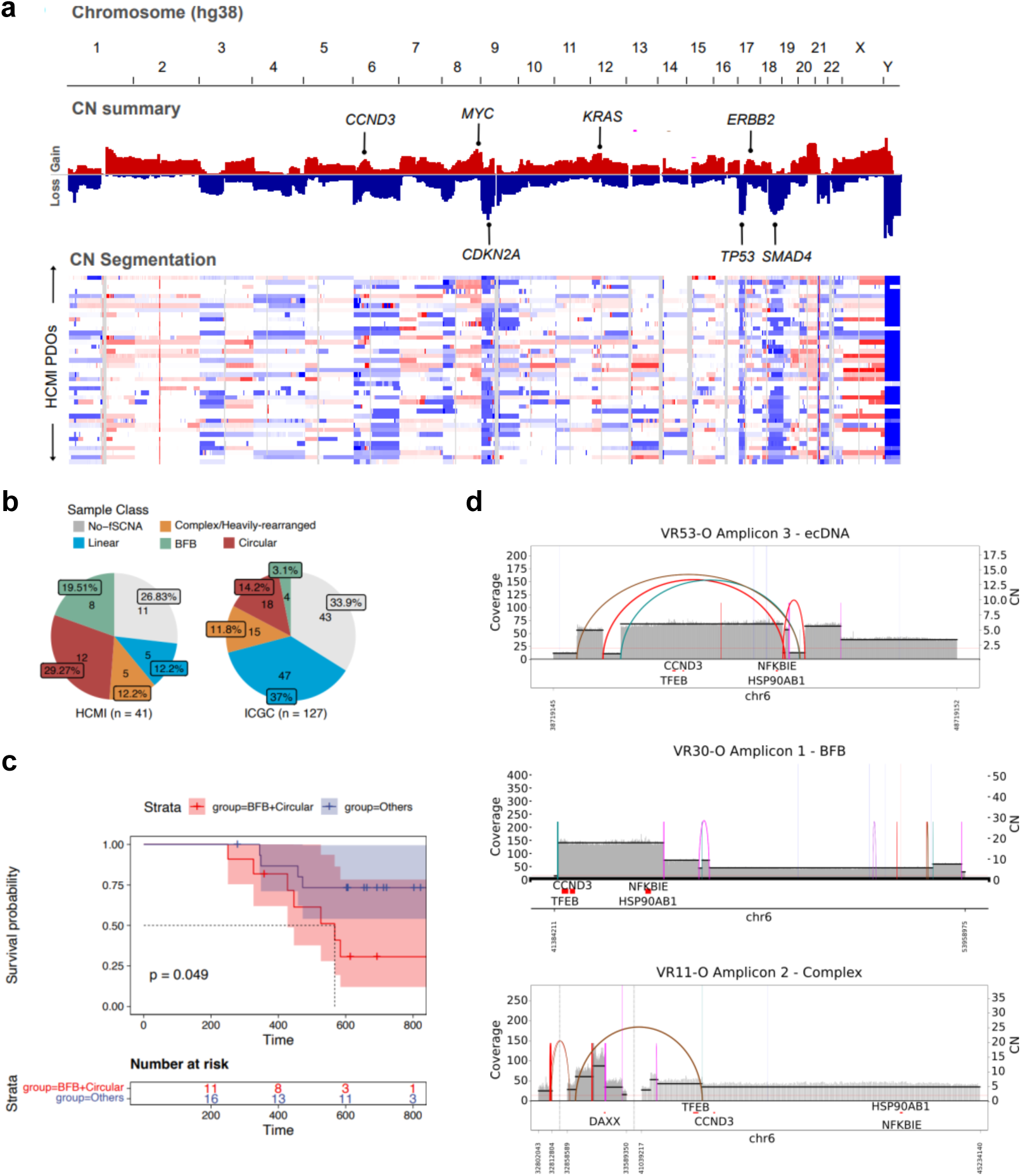
Frequency of ecDNA based amplifications in PDAC. **a**, Copy number (CN) analysis showing: top panel, CN frequency plot displaying the frequency of copy number gains (0.1) and losses (–0.1) observed across the genome (segmentation mean) for the HCMI PDOs (Verona cohort). Representative genes are shown on the plot at their genomic location; bottom panel, CN calls for individual samples. Red represents CN gain and blue represents CN loss. **b**, Pie charts showing proportion of primary tumours (ICGC) and PDOs (HCMI) falling in each sample class based on their existing amplicon types. If a sample contained multiple amplicons, it was classified based on the following order: Circular > BFB > Complex > Linear. If no amplicon was detected, the sample was classified as no-focal somatic copy number amplification detected (No-fSCNA). **c**, Kaplan Meier survival plot comparing the overall survival of patients from the HCMI cohort (n=27) according to AA-base amplicon classification. Survival curves are compared using the log-rank test. **d**, Structural variant (SV) view of AA reconstructed amplicon structures containing the *CCND3* locus for three PDOs with different amplicon classifications. SV view shows coverage depth, copy number segments and discordant genomic connections (curves spanning copy number segments).

**Extended Data Fig 2.**
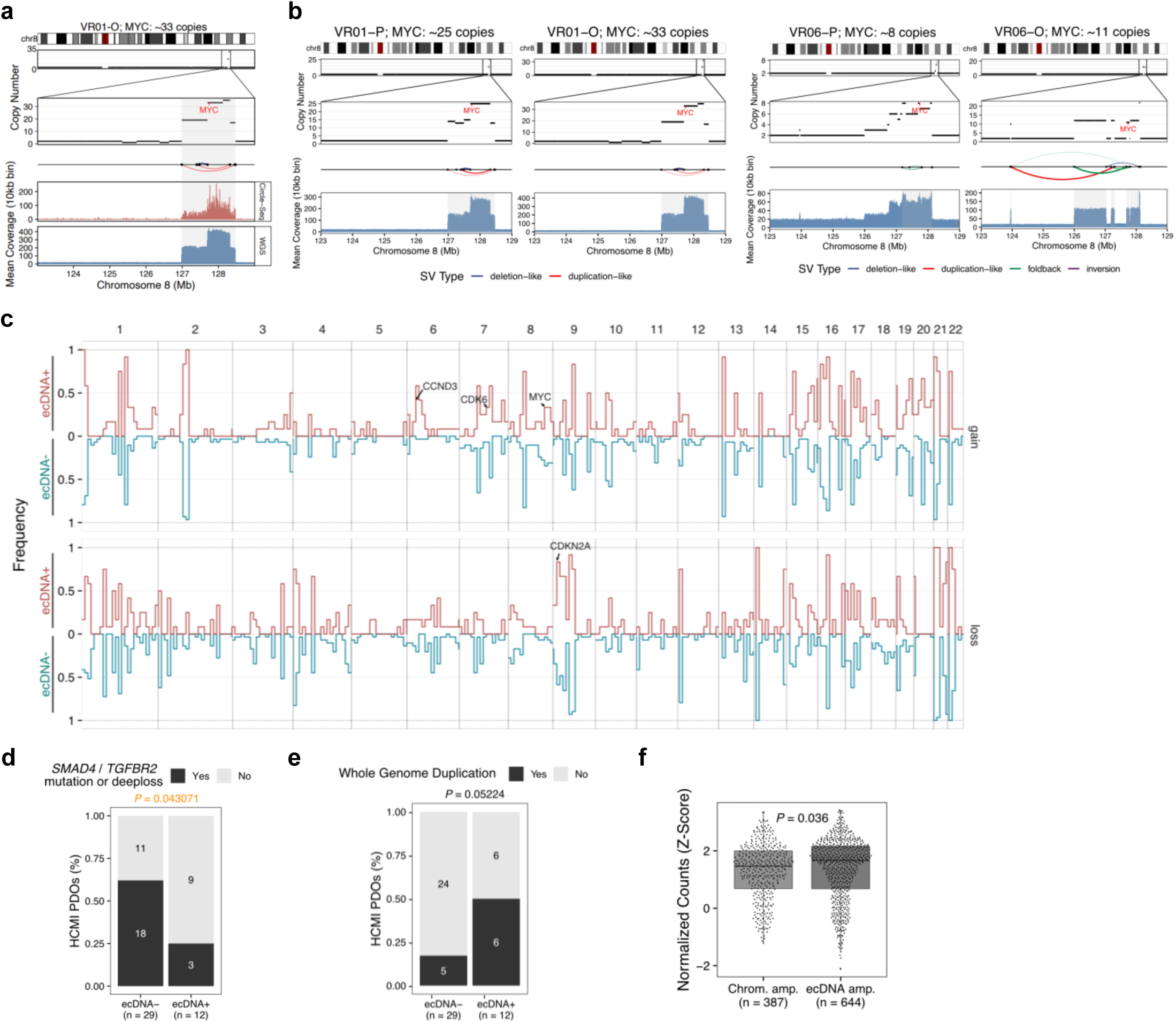
Extrachromosomal *MYC* amplifications are conserved between tissues and paired models. **a**, Validation of the presence of *MYC* on ecDNA by Circle-Seq for VR01-O. The amplified ecDNA segments are highlighted in grey. Mean sequencing coverage was calculated for 10 kb bins. **b**, Copy number alterations on chromosome 8 with a focus on *MYC* region of primary tissues (P) and matched organoids (O) for VR01 (left) and VR06 (right). SVs that connect amplified regions and form ecDNAs are displayed below copy number levels. WGS Coverage is depicted at the bottom. **c**, Genomic overview of copy number alterations. Gain and loss frequency of ecDNA+ (n= 12) and ecDNA-(n = 29) organoids. *CDKN2A* was found to be lost in 10/12 ecDNA+ organoids in comparison to 14/29 ecDNA-organoids (Fisher p value = 0.0026). *CCND3* gain was more common in ecDNA+ organoids (5/12) than ecDNA-organoids (1/29) (Fisher p value = 0.0053) and *CDK6* gain was identified in 4/12 ecDNA+ and 2/29 ecDNA-organoids (Fisher p value = 0.05). Binsize = 10Mbp; Loss: copy number <= 1; Gain: copy number >= 3. **d**, Bar plot showing enrichment for *SMAD4/TGFBR2* inactivating mutations or deep loss in ecDNA-HCMI PDOs. P value was calculated using a two-sided Fisher’s exact test **e**, Bar plot displaying enrichment of whole genome duplication in ecDNA+ HCMI PDOs. P value was calculated using a two-sided Fisher’s exact test. **f**, Boxplot showing normalised expression of genes (Z-scores) located on circular amplicons (ecDNA amp) or chromosomally amplified (chrom amp). Statistical significance was evaluated using a Wilcoxon rank sum test.

**Extended Data Fig 3.**
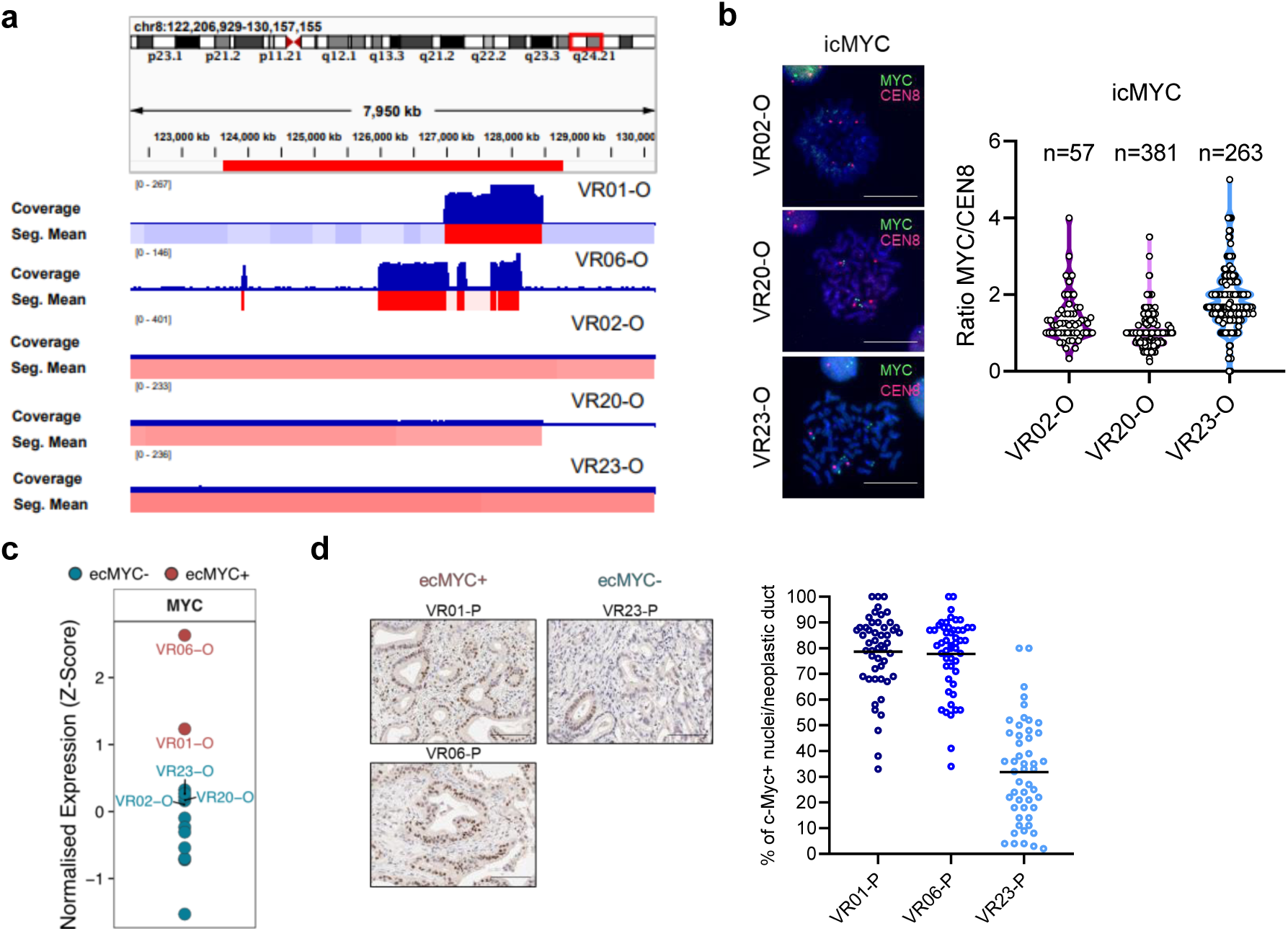
*MYC* copy number heterogeneity in PDAC. **a**, Coverage and segmentation mean histograms spanning the *MYC* locus for the samples indicated. **b**, Representative FISH images of ic*MYC* PDOs metaphases. Scale bar: 20 µm (left). Scattered dot plot showing *MYC/CEN8* ratio for VR02-O, VR20-O, and VR23-O (right). **c**, *MYC* normalised expression values (Z-score) of ec*MYC+* PDOs (red) and ec*MYC*-PDOs (blue). **d**, Representative immunohistochemistry for c-Myc in VR01, VR06, and VR23 patients’ primary tumours. Scale bar: 100 µm (left). Quantification is provided on the right as frequency of c-Myc+ nuclei per neoplastic duct, 50 neoplastic ducts were analysed for each case.

**Extended Data Fig 4.**
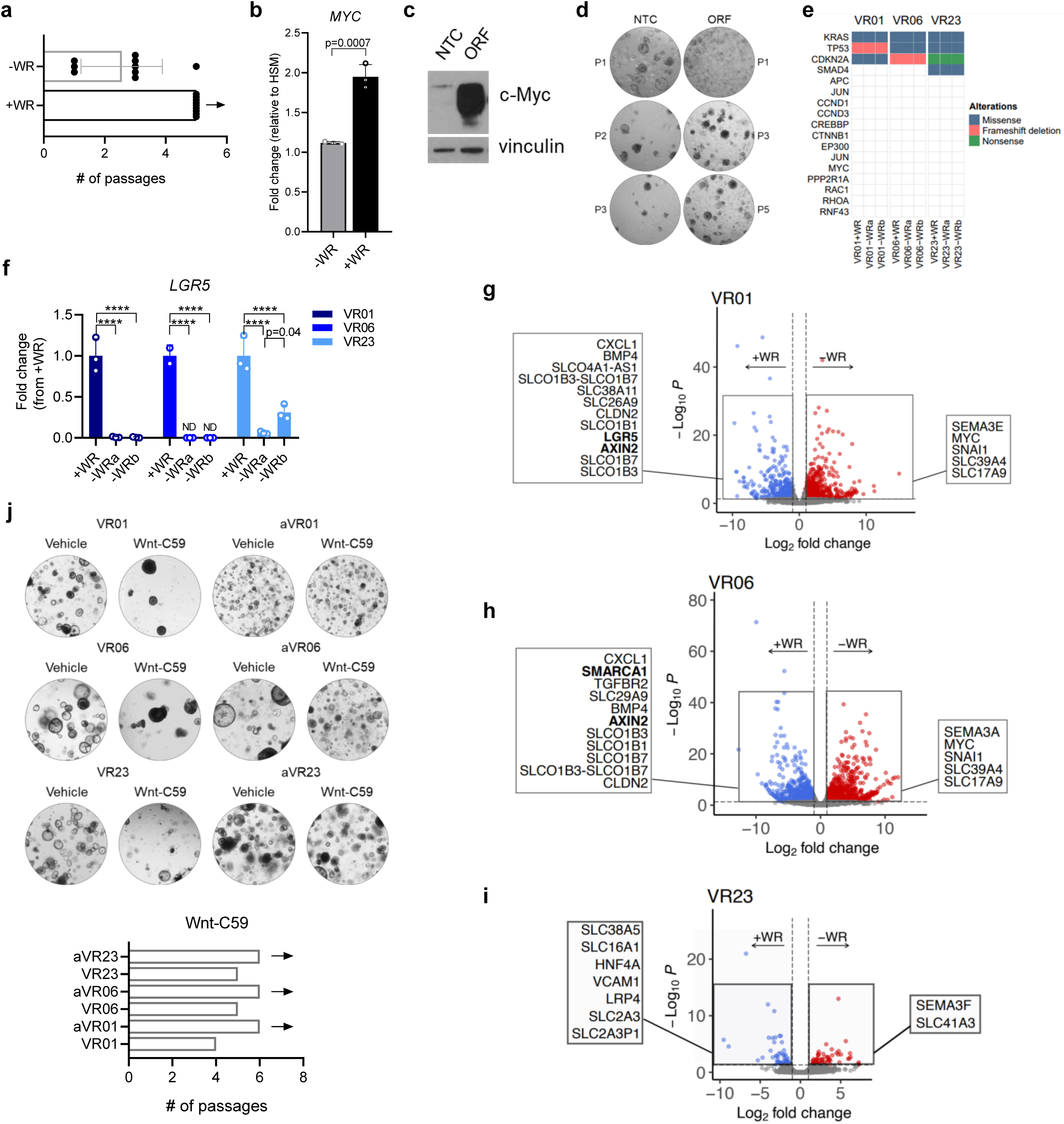
Elevated *MYC* activity enables PDO adaptation to Wnt agonists withdrawal. **a**, Bar plot showing number of passages at which organoid cultures (n = 9) passaged every week with a splitting ratio of 1:3 in –WR media reach extinction, compared to +WR media. **b**, Changes in relative expression levels of *MYC* of starved organoids (HSM) after culture in –WR and +WR media for 8 hours. Results shown as mean ± SD of 3 replicates. P value is reported in the figure as determined by Student’s t-test. *GAPDH* was used as a housekeeping control gene to normalise results. **c**, Immunoblot analysis of c-Myc in whole cell lysate of VR01-O transfected with NTC (non-targeting control) and Myc ORF (open reading frame). Vinculin was used as loading control. **d**, Representative brightfield images of VR01-O transfected with NTC and Myc ORF passaged in –WR media with a splitting ratio of 1:3 every week, showing that VR01-ORF organoids could be propagated in –WR media without extinction of the culture. **e**, Oncoplot displaying absence of mutations in genes involved in Wnt pathway that could explain the acquisition of WR independence of –WR adapted organoids. **f**, Changes in the relative expression levels of *LGR5* in organoids adapted to –WR media compared to baseline (+WR). Results shown as mean ± SD of 3 replicates. Significance was determined by Two-way ANOVA. ****, p<0.0001. *HPRT1* was used as a control. ND, not determined. **g**, Volcano plot showing differentially expressed genes between VR01-O at baseline and after adaptation to grow in –WR media. Upregulated genes were showed as red dots (padj < 0.05 and log2foldchange >1). Downregulated genes were showed as blue dots (padj < 0.05 and log2foldchange <-1). **h**, Volcano plot showing differentially expressed genes between VR06-O at baseline and after adaptation to grow in –WR media. Upregulated genes were showed as red dots (padj < 0.05 and log2foldchange >1). Downregulated genes were showed as blue dots (padj < 0.05 and log2foldchange <-1). **i**, Volcano plot showing differentially expressed genes between VR23-O at baseline and after adaptation to grow in –WR media. Upregulated genes were showed as red dots (padj < 0.05 and log2foldchange >1). Downregulated genes were showed as blue dots (padj < 0.05 and log2foldchange <-1). **j**, Representative brightfield images of baseline (+WR) and adapted organoids (–WR) cultured in the presence of Wnt-C59 (100 nM, PORCN inhibitor) or appropriate vehicle (top). Bar plot showing the number of passages at which each organoid could be propagated in the presence of Wnt-C59 (bottom).

**Extended Data Fig 5.**
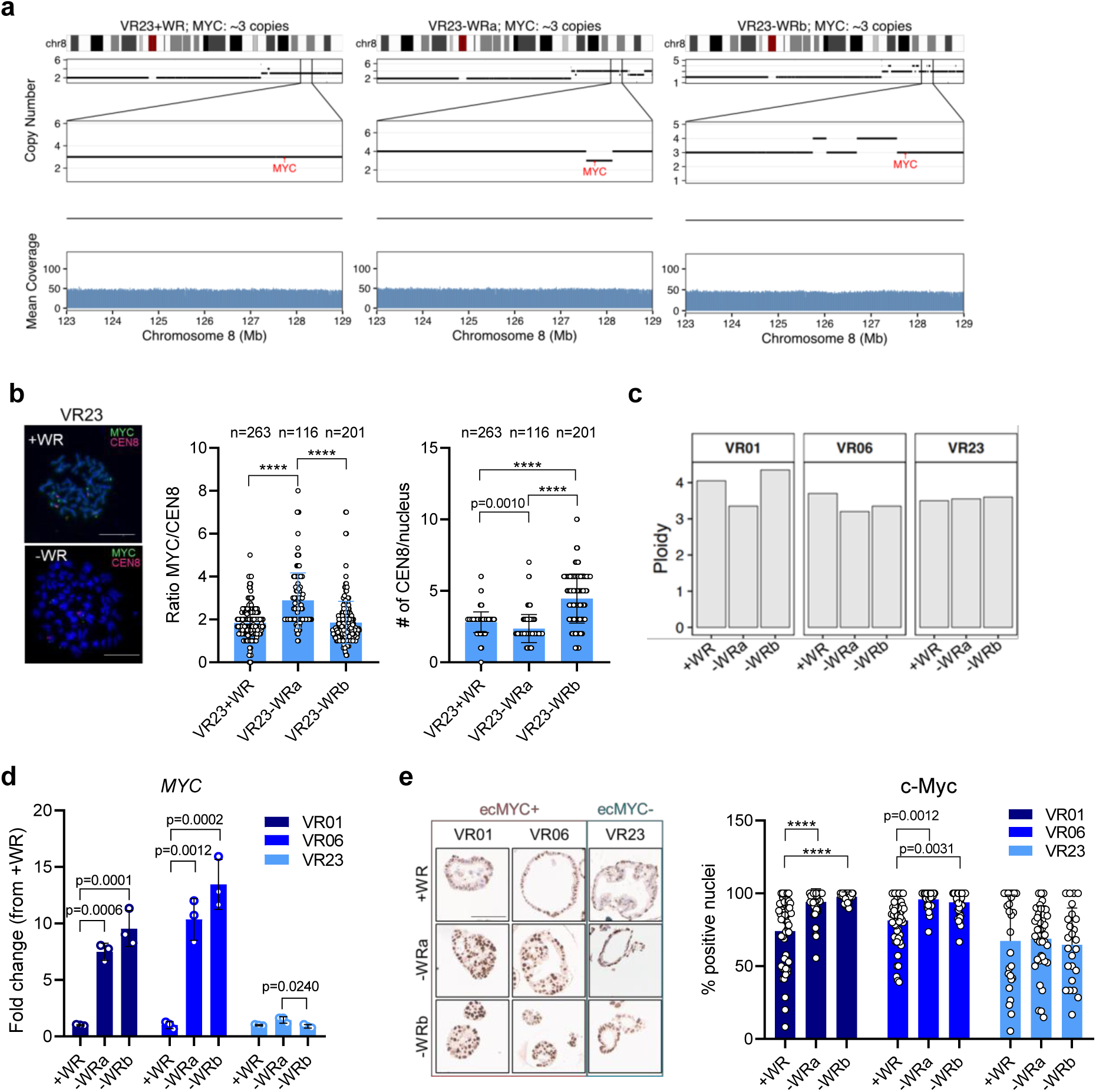
ecDNA supercharges *MYC* expression in adapted PDOs. **a**, Copy number alterations on chromosome 8 with a focus on *MYC* region, of ic*MYC* organoid VR23, at baseline (+WR) and adapted to depleted media (–WR). WGS Coverage is displayed below the copy number level. **b**, Representative FISH metaphases of VR23 at baseline (+WR) and after adaptation to depleted media (–WR). Scale bar: 20 µm (left). Bar plot showing the ratio of *MYC* signal over *CEN8* and the number of *CEN8* spots in VR23 at baseline and after adaptation (–WR, 2 biological replicates) (right). P value by One-way ANOVA. ****, p<0.0001. **c**, Ploidy analysis of organoids at baseline (+WR) and adapted to grow in depleted media (–WR). Ploidy was assessed from the WGS data using AMBER, COBALT, and PURPLE in tumor only mode (https://github.com/hartwigmedical/hmftools). **d**, Changes in the relative expression levels of *MYC* in organoids adapted to depleted media compared to baseline. Results shown as mean ± SD of 3 replicates. P value determined by Two-way ANOVA. ****, p < 0.0001. *GAPDH* was used as housekeeping control gene to normalise results. **e**, Representative immunohistochemistry for c-Myc of formalin-fixed paraffin-embedded organoids at baseline and adapted to grow in –WR media. Scale bar: 100 µm (left). Quantification is provided on the right as frequency of positive nuclei per organoid. A minimum of 25 organoids per sample were analysed. P values determined by Two-way ANOVA. ****, p<0.0001.

**Extended Data Fig 6.**
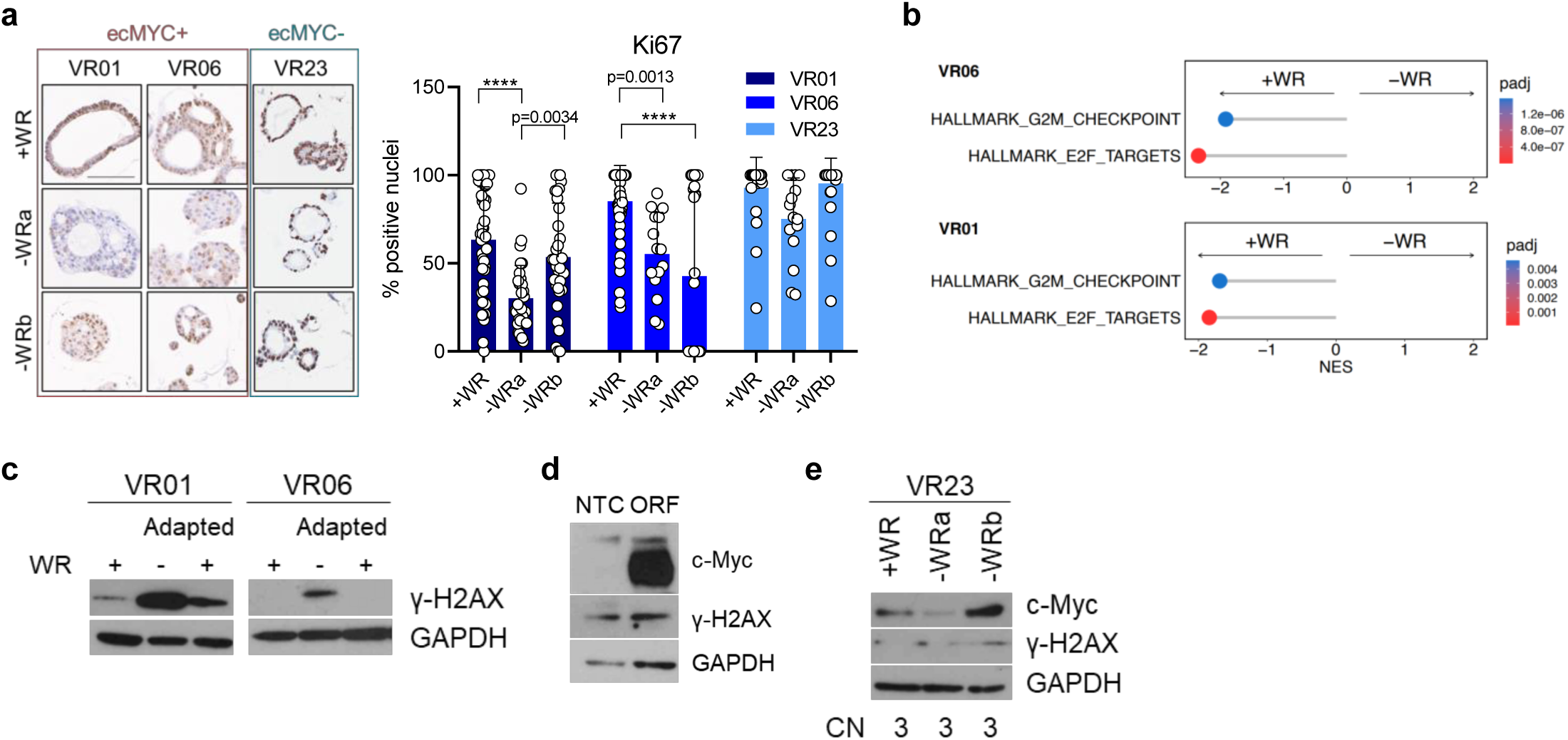
Accumulation of ecDNA leads to increased gH2AX foci. **a**, Representative immunohistochemistry for Ki67 of parental (+WR) and adapted (–WR, 2 biological replicates) organoids. Scale bar: 100 µm. Quantification for Ki67 is provided on the right as frequency of Ki67+ nuclei per organoid, at least 15 organoids were analysed for each condition. Significance was assessed by Two-way ANOVA. ****, p<0.0001. **b**, Enrichment analysis of proliferation-related pathways of ec*MYC* organoids adapted (–WR, NES >0) and at baseline (+WR, NES<0). **c**, Immunoblot of ec*MYC* adapted organoids before and after removal of the imposed pressure. Baseline conditions are included for reference level of proteins expression. GAPDH was used as loading control. **d**, Immunoblot analysis for c-Myc and γ-H2AX in whole cell lysate of VR01-O transfected with NTC (non-targeting control) and *MYC* ORF (open reading frame). GAPDH was used as loading control. **e**, Immunoblot analysis of VR23 at baseline and after adaptation to –WR condition. GAPDH was used as loading control.

**Extended Data Fig 7.**
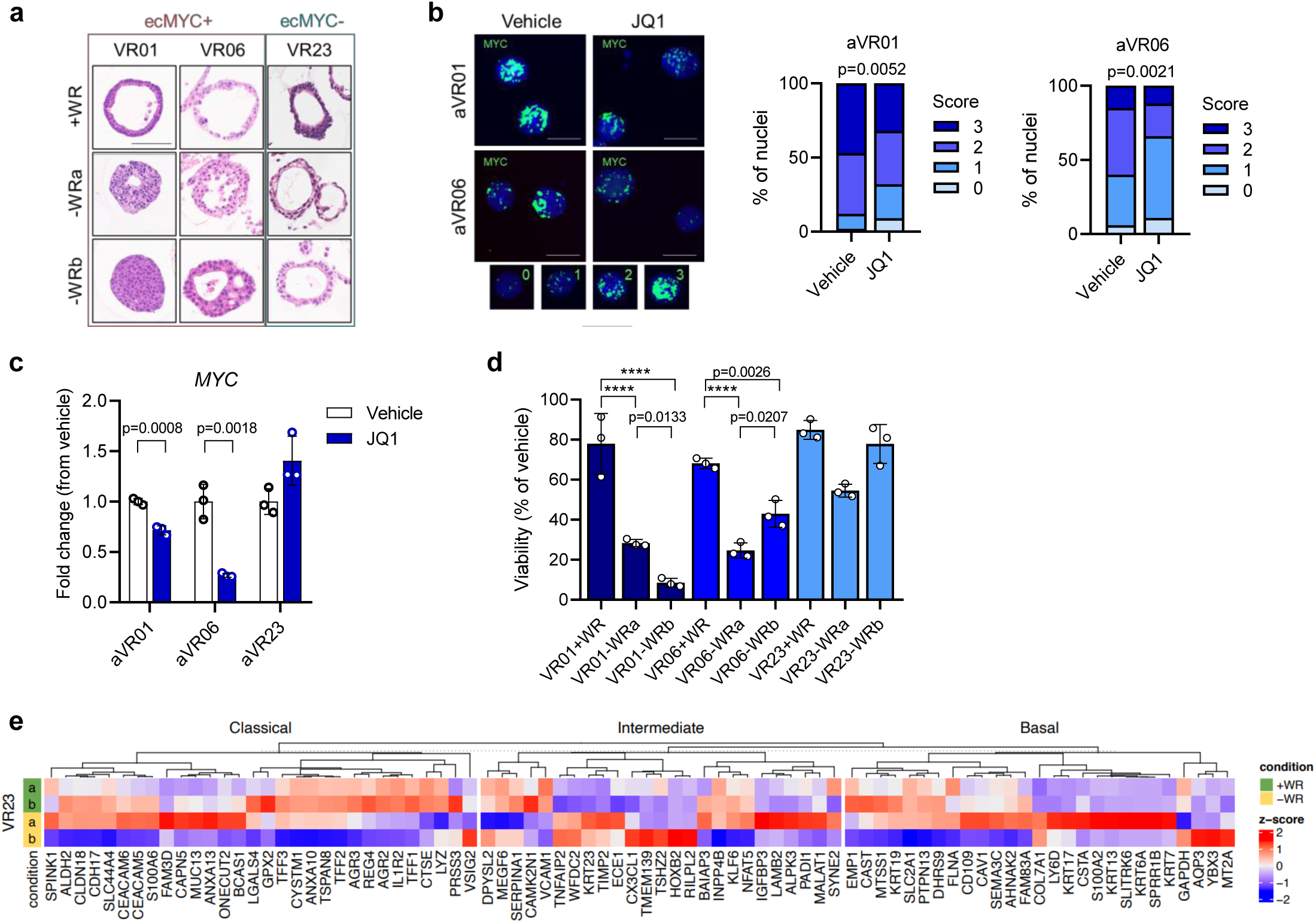
Accumulation of ecDNA is associated with morphological and phenotypic changes. **a**, Representative haematoxylin and eosin (H&E) staining of formalin-fixed paraffin-embedded organoids at baseline and adapted to grow in –WR media. Scale bar: 100 µm. **b**, Representative FISH interphase nuclei of adapted organoids treated with JQ1 (500nM) or appropriate vehicle control for 72 hours showing reduction of *MYC* hubs upon treatment. Scale bar: 20 µm. Representative FISH interphase nuclei for the four different scores (0 to 3) used for the quantification are provided on the bottom. Quantification is provided on the right as frequency of nuclei with different hubs score. Significance was assessed by Chi-square. **c**, Changes in the relative expression levels of *MYC* in adapted organoids treated with JQ1 (500nM) for 72 hours. P value determined by Two-way ANOVA. Results shown as mean ± SD of three replicates. *GAPDH* was used as housekeeping control gene to normalise results. **d**, Bar plot showing cell viability of baseline (+WR) and adapted (–WR) organoids upon 72 hours of JQ1 treatment (500nM). P value determined by Two-way ANOVA. ****, p < 0.0001. **e**, Heatmap displaying the expression of Classical, Intermediate, and Basal genes from Raghavan et al. ^31^ in baseline (2 biological replicas) and adapted (2 biological replicas) VR23 organoids.

## Methods

### Human specimens and clinical data

PDAC tissues were obtained from the General and Pancreatic Surgery Unit at the University of Verona. Written informed consent was obtained from patients preceding the acquisition of the specimens. The fresh tissues used to establish PDOs were collected under a study approved by the Integrated University Hospital Trust (AOUI) Ethics Committee (Comitato Etico Azienda Ospedaliera Universitaria Integrata): approval number 1911 (Prot. n 61413, Prog 1911 on 19/09/2018). Formalin-fixed and paraffin-embedded tissues were collected under protocol number 1885 approved by the AOUI Ethics Committee and retrieved from the ARC-NET Biobank.

## Patient-derived organoid (PDOs) establishment and culture

PDAC PDOs were established following previously published procedures^18^. The specimens used to generate PDOs were examined by pathologists to confirm the presence of neoplastic cells. Briefly, tissue specimens were minced and digested with Collagenase II (5 mg/ml, Gibco) and Dispase I (1.25 mg/ml, Gibco) in human splitting medium (HSM) [Advanced Dulbecco’s Modified Eagles Medium with Nutrient Mixture F-12 Hams (Gibco) supplemented with HEPES (10 mM, Gibco), Glutamax™ (2 mM, Gibco), and Primocin® (1 mg/ml, InvivoGen)] at 37°C for a maximum of two hours, followed by an additional 15-minute digestion with TrypLE (Gibco) at 37°C. The digested material was embedded in Growth factor reduced Matrigel® (Corning) and overlaid with human complete medium (+WR) [Mouse Epidermal Growth Factor (Gibco, 50 ng/ml), B-27 Supplement (Gibco, 1X), Nicotinamide (Sigma-Aldrich, 10 mM), N-Acetylcysteine (Sigma-Aldrich, 1.25 mM), FGF10 (Peprotech, 100 ng/ml), Y-27632 Dihydrochloride (Sigma, 10.5 µM), Gastrin (Tocris, 10 nM), <u>TGFβ Receptor inhibitor A83-01</u> (Tocris, 500 nM), <u>WNT3A Conditioned media</u> (50 % v/v), <u>RSPO1 Conditioned media</u> (10 % v/v), and <u>mNoggin</u> (Peprotech, 100 ng/ml]. Media were refreshed every 3-4 days. For organoid propagation, confluent organoids were removed from Matrigel®, dissociated into small clusters of cells by pipetting, and resuspended in an appropriate volume of fresh Matrigel®. All organoid models were acquired as part of the Human Cancer Model Initiative (HCMI) https://ocg.cancer.gov/programs/HCMI and are available for access from ATCC. The corresponding IDs, along with the clinical data are listed in Supplementary Table 1. Dependency of organoid cultures to WNT3A and RSPO1 was assessed on nine PDOs (VR01, VR02, VR06, VR09, VR20, VR21, VR23, VR29, and VR32). Organoid cultures were passaged once a week with a splitting ratio of 1:3 in +WR or Human Depleted Media (–WR) [Mouse Epidermal Growth Factor (Gibco, 50 ng/ml), B-27 Supplement (Gibco, 1X), Nicotinamide (Sigma-Aldrich, 10 mM), N-Acetylcysteine (Sigma-Aldrich, 1.25 mM), FGF10 (Peprotech, 100 ng/ml), Y-27632 Dihydrochloride (Sigma-Aldrich, 10.5 µM), and Gastrin (Tocris, 10 nM)]. To establish WR independent PDOs, organoids established and propagated in +WR were placed and maintained in –WR for several passages. Due to the cell death induced by –WR, the media was refreshed every three days and Matrigel® every 14 days without propagating the cultures, until the emergence of WR independent PDOs. Growth curve of WR independent PDOs was obtained by plotting the number of domes (one dome refers to 50 µl of Matrigel®) at different days of culture in –WR. Adapted PDOs were reintroduced in +WR or maintained in –WR (control) for five passages before collection of metaphase spreads and proteins. To obtain “Late passage” PDOs, organoids were passaged 40 times post-establishment in +WR medium. For Wnt-C59 experiment, baseline and adapted organoids were passaged every seven days with a splitting ratio of 1:3 in the presence of Wnt-C59 (100 nM, Selleckchem). Wnt-C59 was added to the culture at the day of splitting and after 3 days of culture. Organoids were routinely tested for the presence of *Mycoplasma* contamination using Mycoalert Mycoplasma Detection kit (Lonza).

## Single cells dissociation from organoids

Organoids were incubated with Dispase I diluted in HSM (Dispase I solution, 2 mg/ml) for 20 minutes at 37°C to digest Matrigel®. Following, organoids were dissociated using TrypLE (Gibco) for 10 minutes at 37°C, incubated in Dispase I solution for additional 10 minutes at 37°C, and pipetted to obtain single cells suspension.

## Assessing *MYC* activation by WR media

VR01-O was dissociated into single cells as previously described and plated in Matrigel® in +WR (100,000 viable cells/condition). Following organoids reformation in +WR, PDOs were starved overnight in HSM. Post-starvation, PDOs were stimulated with +WR, –WR, or HSM for eight hours, before collection and isolation of RNA.

## JQ1 *in vitro* treatment

Organoids were dissociated into single cells as previously described. One thousand viable cells were plated in 100 µl 10% Matrigel®/media per well in a 96-well plate in triplicates. JQ1 (500 nM, Selleckchem, S7110) or vehicle were added 40 hours after plating once the organoids were reformed. After 72 hours of treatment, cell viability was assessed using CellTiter-Glo® (Promega) following manufacturer’s instructions. Results were normalised to the vehicle control of each PDO. In parallel, 20,000 viable cells/50 µl Matrigel®, were plated and supplemented with media. Following organoids reformation, cells were treated with JQ1 (500 nM) or vehicle control, and RNA, protein, and metaphase spreads were collected after 72 hours.

## Lentiviral production and infection of organoids

To overexpress *MYC*, we used a lentiviral vector carrying an open-reading frame for *MYC* (mGFP tagged, Origene, cat# RC201611L4). Lentivirus was produced by transfecting the plasmid containing *MYC*, and the packaging plasmid VSV-G with X-tremeGENE9 (Roche, 063665110101) in HEK293T cells. The viral supernatant was harvested 48 hours post-transfection and quantified using Lenti-XTM qRT-PCR Titration kit (Takara Bio) according to manufacturer’s instructions. pLenti-C-Myc-DDK-P2A-Puro lentiORF control particles (Origene, PS100092V) were used as non-targeting control (NTC). For infection, organoids were dissociated into single cells, resuspended in infection media (Dulbecco’s Modified Eagle Medium (DMEM, Gibco), 5 % Foetal Bovine Serum (FBS, Gibco), 1 % Penicillin/Streptomycin (P/S, Gibco)), supplemented with 1 µg/mL polybrene and lentiviral particles (MOI 10). Cells were then spinoculated for one hour at room temperature (RT) and incubated at 37°C for 16 hours. Infected cells were then collected, embedded in Matrigel®, and overlayed with +WR media. Antibiotic selection was started 48 hours after infection using 2 µg/ml puromycin (Gibco).

## Organoids metaphase spreads and interphase nuclei

Organoids were incubated with Colcemid (1 µg/ml, Gibco) in culture media at 37°C and 5 % CO2 overnight. Following incubation, organoids were dissociated into single cells as previously described. Single cells were incubated in hypotonic solution (potassium chloride 0.56 % and sodium citrate 0.8 %) for 20 minutes at RT. Nuclei were then fixed in ice cold methanol-acetic acid (3:1), washed with methanol-acetic acid (2:1), and dropped on adhesion microscope slides.

## DNA Fluorescence *in situ* hybridisation (FISH)

DNA FISH on methanol-acetic acid fixed nuclei was performed using the ZytoLight SPEC MYC/CEN8 Dual Color FISH probe (ZytoVision) while FISH on nuclei from formalin-fixed paraffin-embedded tissues and organoids was performed using a Vysis LSI MYC Break Apart Rearrangement Probe kit (Abbott). Before hybridisation, tissues were deparaffinised and rehydrated, pre-treated with 0.1 citrate buffer (pH 6) solution at 85°C for 30 minutes, followed by pepsin treatment (4 mg/ml in 0.9% NaCl, pH 1.5) for four minutes at 37°C. For both tissues and PDOs, the probes were applied to the slides and sealed with rubber cement and incubated in a humidified atmosphere (Thermobrite System) at 80°C for 10 minutes to allow denaturation of the probes and of the DNA target. Slides were then incubated overnight at 37°C to allow for hybridisation. The rubber cement and the coverslip were then removed, and the slides were washed in 2X SSC/0,3% NP40 for 15 minutes at RT and then at 72°C for two minutes. Following post-hybridisation washes, slides were counterstained with DAPI 1 ug/ml (Kreatech, Leica).

For tissues and embedded organoids, images were acquired on Leica DM4B Fluorescent microscope. *MYC*-targeting probe was acquired in red (Rhodamine) and pseudo-coloured as green to match *MYC* signal from PDOs. Nuclei were acquired and visualised in blue (DAPI).

For PDOs, images were acquired either on Leica DM4B, ZEISS Axio Imager 2 or Leica TCS SP5 Fluorescent microscopes. The *MYC*-targeting probe was acquired and shown in green (L5 for Leica, GFP for ZEISS). For the Leica microscopes, the *CEN8*-targeting probe was acquired and shown in red (Rhodamine). For ZEISS, the *CEN8*-targeting probe was acquired in Orange (DsRed) and pseudo-coloured as red to match the *CEN8* signal from Leica. Nuclei were acquired and visualised in blue (DAPI). Number of fluorescent signals for each probe for each nucleus, for both tissues and PDOs, was quantified with FIJI (ImageJ2 version 2.9.0/1.53t). Number of ecDNA+ and HSR+ metaphases were counted by visual inspection of slides. To quantify hubs of overlapping signal, a scoring system (0-3) was devised, where each score corresponded to a level of overlapping signal. Representative images for each score are shown in FigureS7b.

## Histology and immunostaining

For histopathological analysis, organoids were released from Matrigel® using Dispase I solution as previously described, fixed with 10% neutral buffered formalin for 20 minutes, and embedded in Histogel Processing Gel (FisherScientific). Histogel-embedded organoids were processed according to routine histology procedures and embedded in paraffin. To account for effect of the media, +WR PDOs were put in –WR for 24 hours prior to embedding and fixation. Haematoxylin and Eosin (H&E) and immunostainings were performed on sections of formalin-fixed, paraffin-embedded tissues and organoids, following established procedures using the reported primary antibodies: c-Myc (Abcam, cat# ab32072), GATA6 (R&D Systems, cat#AF1700), ΔNp63 (Leica, clone BC28, cat#PA0163), CK5 (Novocastra, clone XM26, cat# PA0468), Ki67 (Abcam, cat#ab16667), and γ-H2AX (eBioscience, clone CR55T33, cat#14-9865-82). Immunohistochemistry slides were then scanned and digitalised using the Aperio Scan-Scope XT Slide Scanner (Aperio Technologies). In tissues, c-Myc staining was quantified as a percentage of positive nuclei per neoplastic duct using Aperio ImageScope. In organoids, c-Myc, Ki67, and GATA6 staining were quantified as percentage of positive nuclei per organoid, using Aperio ImageScope. For immunofluorescence, images were acquired by Leica TCS SP5 Fluorescent microscope and quantify using ImageJ (https://imagej.nih.gov/).

## Immunoblotting

Proteins were prepared using Cell Lysis Buffer (Cell Signaling Technology) supplemented with Protease inhibitor cocktail (Sigma) and Phosphatase inhibitor PhosphoSTOP (Roche). Protein lysates were separated on 4-12 % Bis-Tris NuPAGE gels (Life technologies), transferred to a PDVF membrane (Millipore) and then incubated with the reported antibodies: c-Myc (Abcam, cat# ab32072), γ-H2AX (Abcam, cat# ab81299), Cleaved PARP (Cell Signaling Technologies, cat#9541), GAPDH (Cell Signaling Technologies, cat# 5174), and vinculin (Cell Signaling Technologies, cat# 4650). Quantification of immunoblots bands was performed using ImageJ. To account for effect of the media, +WR PDOs were put in –WR for 24 hours prior to collection of the cells’ pellet.

## Quantitative real time polymerase chain reaction (qRT-PCR) analysis

RNA from organoids were isolated using the TRIzol® Reagent (Life Technologies), followed by the column based PureLink RNA Mini Kit (Thermo Fisher Scientific). Reverse transcription of 1 µg of RNA was performed using the TaqMan® Reverse Transcription reagents (Applied Biosystems), and 20 ng of cDNA was used in the PCR reaction. The following TaqMan® probes *HPRT1* (Hs02800695_m1) and *LGR5* (Hs00173664_m1) were used in the project. The following primers (Eurofins) were used with SYBR™ Green PCR master mix (ThermoFisher):

*MYC* Forward: CCTGGTGCTCCATGAGGAGAG

*MYC* Reverse: CAGACTCTGACCTTTTGCCAG

*GAPDH* Forward: ACAGTTGCCATGTAGACC

*GAPDH* Reverse: TTTTTGGTTGAGCACAGG

Relative gene expression quantification was performed using the ΔΔCt method with the Sequence Detection Systems Software, Version 1.9.1 (Applied Biosystems).

## DNA Isolation

Organoids were incubated in Cell Recovery Solution (Corning) for 30 minutes at 4°C to remove Matrigel®, and were pelleted by centrifuging 10,000 g for 5 minutes at 4°C. For tissues, slices from snap frozen PDAC tissues were assessed by a pathologist for percent neoplastic cellularity and only tissues with higher than 20 % neoplastic cellularity were used. For WGS and panel DNA sequencing, DNA isolation was performed using DNeasy Blood & Tissue kit (Qiagen). For CIRCLE-Seq, high molecular weight DNA was extracted using the MagAttract HMW DNA Kit (Qiagen).

## Whole genome sequencing (WGS)

DNA quality was assessed by DNF-467 Genomic DNA 50 kb Kit on a Bioanalyzer 2100 (Agilent). Libraries were prepared and sequenced using NovaSeq 6000 S4 Reagent Kit v1.5 (300 cycles) at 15x coverage 160 million reads per sample.

### Data pre-processing and alignment

Sequencing data were pre-processed and mapped to the reference genome using the nf-core/sarek pipeline (version 3.0.2) ^46^. In short, Fastp (version 0.23.2) ^47^ removed low-quality bases and adapters, BWA Mem (version 0.7.17-r1188)^48^ mapped trimmed reads to the reference genome GRCh38, provided by the Genome Reference Consortium (https://www.ncbi.nlm.nih.gov/grc), mapped reads were marked for duplicates using Picard Markduplicates, and read base quality scores were recalibrated using GATK BaseRecalibrator and GATK ApplyBQSR ^49^.

### Amplicon Characterisation

The nf-core/circdna (version 1.0.1, https://github.com/nf-core/circdna) pipeline branch ‘AmpliconArchitect’ was used to define amplicon classes in each WGS sample. Nf-core/circdna calls copy number using cnvkit (version 0.9.9) ^50^ and prepares amplified segments with a copy number greater than 4.5 for AmpliconArchitect by utilising functionality of the AmpliconSuite-Pipeline (https://github.com/jluebeck/AmpliconSuite-pipeline). AmpliconArchitect (version 1.3_r1) ^21^ was ran on the aligned reads and the amplified seeds to delineate the amplicon structures. Identified amplicons were then classified using AmpliconClassifier (version v.0.4.11) ^51^ into circular (ecDNA), linear (linear amplicon), complex (complex amplicon), or BFB (amplicon with a breakage-fusion-bridge signature). Samples containing at least one circular amplicon (ecDNA) were termed “ecDNA+”, whereas samples without ecDNA amplicons were termed “ecDNA-”. Samples were also classified into ‘Circular’, ‘Linear’, ‘Complex’, ‘BFB’, or ‘no-fSCNA’ (no-focal somatic copy number amplification detected) by the types of amplicons they contained (see Kim et al. ^22^. Samples with multiple amplicons were classified based on the amplicon with the highest priority. The priority is: Circular > BFB > Complex > Linear.

### Copy number calling

Copy number calls of the WGS samples were generated by cnvkit (version 0.9.9) ^50^. The identified segments were then classified as gain (copy number >= 3), loss (copy number <= 1), or deeploss (copy number <=0.25).

### Chromosomal instability signatures

Chromosomal instability signatures, including the CX9 replication stress signature, were assessed from the WGS copy number profiles using the R-package CINSignatureQuantification ^26^.

### Ploidy analysis

Sample ploidy was derived using PURPLE ^52^, which estimates copy number and ploidy by using read depth ratio and tumour-B-allele frequency (BAF) from COBALT and AMBER, respectively (https://github.com/hartwigmedical/hmftools). COBALT, AMBER, and PURPLE were used in tumour-only mode using their default parameters. Notably, PURPLE was used with a fixed parameter value of purity set to 1 for all samples, ensuring consistency in the analysis.

## Circularisation for *in vitro* reporting of cleavage effects by sequencing (CIRCLE-seq)

To enrich circular DNA for sequencing, each DNA sample was digested for seven consecutive days with ATP-dependent Plasmid-Safe DNase (Lucigen) to remove linear/chromosomal DNA. Each day 20 units of enzyme and 4 µl of a 25 mM ATP solution were added. After seven days, the DNase was heat-inactivated for 30 minutes at 70°C. The fold change reduction in linear DNA was assessed by qPCR targeting the chromosomal gene *HBB* and the mitochondrial gene *MT-CO1*. Amplification of circular DNA was performed with a Phi29 polymerase as described in Koche et al. ^53^. Amplified circular DNAs were then prepared for sequencing. In short, around 550 ng of DNA were sheared to a mean length of around 400-450 base pairs (bp) and subjected to library preparation using the NEBNext Ultra II DNA Library Prep Kit for illumina (NEB), which included sequencing adapter addition, and amplification. DNA Clean-up was performed using the Agencourt AMPure XP magnetic beads. All prepared libraries were sequenced using the Illumina NextSeq500 with the NextSeq 500/550 Mid Output Kit v2.5 (300 Cycles), generating around 10M paired end 150bp reads per sample.

### Data processing

Sequencing reads were trimmed for both quality and adaptor sequences using cutadapt (version 3.4)^54^. Trimmed reads were aligned to the GRCh38 reference genome using BWA Mem (version 0.7.17-r1188) ^48^.

### Identification of sequencing coverage

Sequencing read coverage per 50 bp bin was calculated using deeptools ‘bamCoverage’ (version 3.5.1^55^) with default values. For visualisation, the 50 bp read coverage values were combined into 10,000 bp bins using the function ‘ScoreMatrixBin’ of the genomation (version 1.2.6) R-package^56^.

## DNA Panel Sequencing

Library preparation was performed using SureSelectXT HS Target Enrichment System (Agilent). Panel pair-end 2×150 sequencing was performed on NextSeq 550 (Illumina). Genes present in the panel are reported in Supplementary Table 3.

## RNA Sequencing (RNA-seq)

RNA from organoids were isolated using the TRIzol® Reagent (Life Technologies), followed by the column based PureLink RNA Mini Kit (Thermo Fisher Scientific). Purified RNA quality was evaluated using RNA 6000 Nano kit on a Bioanalyzer 2100 (Agilent) and only RNA with an RNA Integrity Number (RIN) greater than 9 were used. RNA-seq library were obtained using poly(A) enrichment with TrueSeq Stranded mRNA Library Prep kit (Illumina). Libraries obtained from PDOs at baseline (n = 14, analyses displayed in Figure 1) were sequenced to a depth of 30M fragments and 150 base paired end reads on an Illumina NextSeq 500 sequencer. For comparison between +WR and adapted to –WR PDOs, +WR PDOs were put in –WR for 24 hours prior to RNA collection, to account for effect of the media. The resulting libraries were sequenced to a depth of 11M fragments for organoids and 75 base paired end reads on an Illumina NextSeq 500 sequencer.

## RNA-Seq Analysis

For downstream analyses, the raw counts were normalised using the ‘rlog’ function of the DESeq2 R-package. Genes with less than a total of 20 counts across all PDOs were removed prior normalisation. To compare gene expression values across amplicon types, the normalised gene values were Z-score normalised.

### Gene Set Enrichment Analysis

Differential gene expression analysis was conducted using ‘DESeq2’ ^57^. Log2 fold change shrinkage was applied using the ‘lfcShrink’ function in ‘DESeq2’ with the ‘ashr’ method ^58^. Gene set enrichment analysis (GSEA) was performed using the ‘fgsea’ R-package ^59^ with the Hallmark pathways database provided by the ‘msigdbr’ R-package ^60^

### Subtyping

The subtyping was performed scoring the samples according to the Raghavan signatures ^31^ with the gsva function and assigning the subtype according to what signature (basal or classical) achieved the highest score.

### Fusion analysis

Fusion analysis was performed on adapted organoids to exclude the presence of chimeric proteins reactivating the wnt pathway. The nf-core/rnafusion pipeline was used to evaluate gene fusion from our RNAseq data; the pipeline was run under default parameters using all the fusion detection tools provided (arriba, fusioncatcher, pizzly, squid, starfusion and stringtie). Only fusions detected by at least two tools were considered as confident.

## Survival analysis

Patient survival data was utilised for Kaplan-Meier survival analysis. This data encompassed the patients’ vital status, the number of days to death or to the last follow-up point. The analysis was performed using the R-packages ‘survival’ (v3.4-0) and ‘survminer’ (v0.4.9). Patients who died due to ‘Surgical complications’ or ‘Infection’ were excluded from the analysis

## Statistical analyses

All statistical analyses were carried out using R (v4.1.2) or GraphPadPrism (v9.5.1). A Fisher’s exact test and Chi Square were used to evaluate the significance in contingency tables. The Wilcoxon rank-sum test was used in two-group comparisons and the relationship between two quantitative variables was measured using the Pearson correlation. Other statistical tests performed are described in the figures or in the figure legends.

## Public Datasets

Amplicon information for the International Cancer Genome Consortium (ICGC) PACA-CA and PACA-AU whole-genome sequencing (WGS) samples was obtained from ^22^. Additional matching ploidy data was retrieved from the ICGC Data Portal (rhttps://dcc.icgc.org/releases/PCAWG). To focus on PDAC specifically, only PDAC tumours with histological types ‘8500/3’, ‘8560/3’, ‘8140/3’, ‘Adenosquamous carcinoma’ and ‘Pancreatic Ductal Adenocarcinoma’ were used in the downstream analysis.

## Acknowledgement

We gratefully acknowledge the Centro Piattaforme Tecnologiche (CPT – University of Verona, Verona, Italy) for granting access to the genomic and imaging facilities of the University of Verona.

## Authors’ contributions

AM, DS, EF, PB and VC conceived and designed the research; SDa and SA established human organoids cultures; AM and EF performed experiments with organoids; AM, EF and SP performed FISH experiments; AM, EF, MB, FL, and LV performed the immunostaining experiments. DS, DP and PB analysed omics data and generated displays; CN assisted with the circle-seq analysis. RTL, RS, GM, AR, AG, MM and SG collected samples, provided tissue annotation and clinicopathological information; AS and CL performed histopathological evaluation of human tissues. AM, DS, EF, PB, and VC interpreted the data with assistance from DAT and CP; AM, DS, EF, PB and VC wrote the manuscript. VC and PB supervised the study. All authors approved the final version of the manuscript.

## Supplementary information

Supplementary Table 1_This table contains clinical data and histology of tumours used to established HCMI patient-derived organoids (Verona’s cohort).

Supplementary Table 2_This table contains focal amplification classifications of organoids with details about interest chromosomal segments.

Supplementary Table 3_This table contains the list of genes included in the panel and the high coverage targeted sequencing results.

Supplementary Table 4_This table contains differential gene expression and pathway enrichment analysis of ecDNA+ compared to ecDNA-samples.

Supplementary Table 5_This table contains differential gene expression and pathway enrichment analysis of adapted organoids compared to baseline condition.

